# Shu complex is an ATPase that regulates Rad51 filaments in homologous recombination-directed DNA damage response

**DOI:** 10.1101/2024.04.22.590616

**Authors:** Sam S. H. Chu, Guangxin Xing, Vikash K. Jha, Hong Ling

## Abstract

Rad51 filaments are Rad51-coated single-stranded DNA and essential intermediates in homologous recombination (HR) and the HR-associated DNA damage response. The yeast Shu complex (Shu) is a conserved regulator of HR, working through its modulation of Rad51 filaments. However, the biochemical properties of Shu remain unclear, which hinders molecular insight into Shu’s role in HR and the DNA damage response. In this work, we biochemically characterized Shu and analyzed its molecular actions on single-stranded DNA and Rad51 filaments. First, we revealed that Shu preferentially binds DNA with ssDNA components and ssDNA/double-stranded DNA junctions. Then, we identified and validated, through site-specific mutagenesis, that Shu is an ATPase and hydrolyzes ATP in a DNA-dependent manner. Furthermore, we showed that Shu interacts with ssDNA and Rad51 filaments at the 5’ end preferentially, altering the conformations of ssDNA and the filaments. The alterations depend on the ATP hydrolysis of Shu, suggesting that the ATPase activity of Shu is important in regulating its functions in HR. The preference of Shu for acting on the 5’ end of Rad51 filaments aligns with the observation that Shu promotes lesion bypass at the lagging strand of a replication fork. Our work on Shu, a prototype modulator of Rad51 filaments in eukaryotes, provides a general molecular mechanism for Rad51-mediated error-free DNA lesion bypass.

## Introduction

Living organisms are constantly subjected to endogenous and environmental factors that cause DNA damage. About 10^4^-10^5^ DNA lesions appear per mammalian cell per day due to DNA damage(1). Cells deploy DNA repair and DNA damage tolerance pathways to maintain genomic integrity for both cell survival and accurate transmission of genetic information to the next generation(2, 3). Operating throughout the entire cell cycle, DNA repair pathways recognize and excise damaged sites, and restore the original sequence by subsequent DNA synthesis with high fidelity(1, 4). In contrast, DNA replication-blocking lesions that persist to S and G_2_ cell cycle phases are bypassed by DNA damage tolerance(5, 6). Stalling of DNA replication forks by lesions inevitably leads to their collapse, resulting in cell death. Hence, cells use mechanisms for tolerance of DNA damage to bypass the lesion sites and finish DNA synthesis without actually removing the lesions(2, 7).

There are two types of DNA damage tolerance mechanisms: error-prone translesion synthesis (TLS) and error-free DNA lesion bypass(2, 8). In TLS, specialized DNA polymerases accommodate structurally distorted and bulky DNA lesions to directly replicate across them in DNA(9–11). However, TLS has increased mutation rates due to loose active sites and lack of proofreading activity in TLS polymerases(11). In contrast, the error-free DNA lesion bypass, mediated by homologous recombination (HR), does not cause additional mutations(12). The highly conserved Rad51 recombinase protein and its paralogs play essential roles in error-free lesion bypass. Rad51 binds single-stranded DNA (ssDNA) to form stable filaments for HR actions(5, 12). The hallmark of the error-free pathway is to use homologous sister chromatids as the templates to ensure faithful replication(5, 12). Rad51 filaments mediate essential steps associated with template switching to ensure genomic integrity. They cooperate with various Rad51 paralogs to facilitate homology searching, strand invasion and strand exchange with the homologous sister chromatids, leading to the eventual bypass of the DNA lesion(5, 12, 13).

A yeast Rad51 modulator protein has been identified as the Shu complex (Shu) in error-free DNA lesion bypass. This complex is coded by four *shu* genes, *csm2, psy3, shu1 and shu2,* which belong to one epistasis group(14, 15). The Shu complex is highly conserved from yeast to humans(16–20). The *shu* gene defects have been linked to various genetic disorders and cancers in humans(21, 22). In *Saccharomyces cerevisiae* (budding yeast), mutations in the *shu* gene have been associated with DNA damage sensitivity to methyl methanesulfonate (MMS) and hydroxyurea (HU), which cause genomic instability(13, 15, 23). The gene products, Csm2, Psy3, Shu1 and Shu2, form the Shu complex heterotetramer(24, 25). Structural studies revealed that Csm2, Psy3 and Shu1 share structural similarities with Rad51’s ATPase core domain(26–28). Thus, the Shu proteins are considered to be Rad51 paralogs(5, 26–28). Shu proteins structurally mimic Rad51 and stabilize Rad51 filaments(5, 26, 27, 29), in turn, promote error-free DNA lesion bypass(29). Specifically, the Csm2-Psy3 dimer in the Shu complex has been shown to facilitate Rad51-ssDNA filament formation at stalled replication forks(30). Recently, studies from the Bernstein group further demonstrated that the Shu complex directly promotes lesion-specific HR processes to avoid double-stranded (ds) breaks and reduces toxicity of alkylation damage of DNA(20, 31). These findings provide clear evidence that the Shu complex plays an important role in error-free lesion tolerance.

Biochemically, the Csm2-Psy3 dimer from the Shu complex has been shown to preferentially bind either dsDNA forks or 3’-overhangs, which are common DNA structures during HR(32). ATPase activity has been reported in the human Shu homolog(16), and a remodeling function on Rad51-ssDNA filaments has been shown in the *C*. *elegans* homolog (18, 33). However, whether the prototype yeast homolog, the Shu complex, has similar biochemical functions and how these biochemical functions might direct their roles in the DNA damage response remain unknown.

In this study, we report biochemical characterization of the Shu complex to provide molecular insights on its role in error-free lesion bypass. We demonstrated that the Shu complex binds DNA with ssDNA components preferentially. We found that the Shu complex binds nucleotides (ATP/ADP) through an ATP-binding domain that closely resembles Rad51 Walker A motif in structure. Furthermore, we revealed that the Shu complex possesses DNA-stimulated ATPase activity and confirmed the enzymatic activity by site-specific mutagenesis. Lastly, we showed that the Shu complex changes the conformations of ssDNA and Rad51-ssDNA filaments. The conformational change is primarily dependent on the ATP hydrolysis activity of the Shu complex. Taken together, our work reveals important biochemical properties of the Shu complex and its ability to physically alter Rad51 filament conformation. This work helps us understand the molecular mechanism of error-free lesion bypass.

## Materials and Methods

### Molecular cloning of proteins

The plasmids pET-Duet-Shu2-Shu1 encoding 6xHis-Shu2 and native Shu1, and pCDF-Duet-Csm2-Psy3 encoding native Csm2 and Psy3 with S-tag at the C-terminus were kindly provided by Dr. Wei Xiao of University of Saskatchewan(23). We made all the constructs with the polymerase incomplete primer extension (PIPE) method(34). SHU2 and SHU1 were cloned into the modified pET-Duet vector mixed with a plasmid pMCSG9(35) component for expression of Shu2 with a His-MBP tag and native Shu1. CSM2 and PSY3 were similarly cloned for expression of Psy3 with a His-MBP tag and native Csm2. The pCDF-Duet-Csm2-Psy3 plasmid used for co-expression with Shu2-Shu1 was modified to express a native, tag-free Csm2-Psy3 dimer. The mutants K199A/R200A/R201A of Psy3 and K189A/R190A/R191A/R192A of Csm2 for DNA-binding deficiency, and K52A (Walker A) and D136A (Walker B) of Psy3 were made for function validation. Yeast RAD51 was cloned from *Saccharomyces cerevisiae* genomic DNA and inserted into a pMCSG7 derivative with a His-MOCR tag(37).

### Protein expression and purification

The expression plasmids were transformed into the *Escherichia coli* BL21 (DE3) pRARE strain for overexpression. For the complete Shu complex, plasmids containing Shu2-Shu1 and Csm2-Psy3 were transformed together for co-expression. Transformed cells were grown in LB medium at 37 °C containing appropriate antibiotics. The LB media was supplemented with 10 mM final concentration of phosphate to inhibit alkaline phosphatase, a common ATPase contaminant expressed in *E*. *coli*. Cell cultures were grown until OD_600_ reached 0.6 to 0.8 and then induced with 0.25 mM IPTG and 50 μM ZnCl_2_ at 16 °C for 16 hours. Cultures were harvested by centrifugation and stored at −80 °C until used in purification.

Cell pastes were lysed in a lysis buffer (50 mM Tris-HCl, pH 7.5, 500 mM NaCl, 5 % glycerol, 1 mM PMSF, 1 mM Benzamidine and 5 mM βMe). Clear cell lysates (clarified by centrifugation) were purified with Ni-HiTrap FF and MBPtrap HP columns (GE Healthcare). Then tag-cleaved proteins was purified with ion exchange chromatography (HiTrap SP FF column (GE Healthcare). Target proteins in the HiTrap SP FF column were eluted by a linear gradient of NaCl and further purified by gel filtration chromatography using a Superdex 200 10/300 GL column (GE Healthcare). The purified proteins were of high purity (>95 %) and homogeneity as determined by Bis-Tris-SDS-PAGE. Protein concentration measurements were calibrated using amino acid analysis.

### Electrophoretic Mobility Shift Assays (EMSAs) for DNA binding

DNA-binding reactions (20 μl) were carried out for 30 min at 4 °C in binding buffer (50 mM Tris-HCl, pH 7.5, 50 mM NaCl, 5 mM MgCl_2_ and 2 mM DTT) with the increasing concentrations of Shu complex and 0.2 μM DNA substrate. The protein-DNA mixtures (15 μl each) were resolved in 5 % native polyacrylamide gels with 15 % bottom layer (to capture unbound DNA substrates). All assays were repeated at least three times.

### Fluorescence Polarization Assays (FPAs) for DNA binding

DNA binding assays were conducted as described (27, 32). Experiments were performed using a VICTOR^3^ V 1420 Multilabel Counter (Perkin Elmer). Fluorescence polarization measurements were recorded using the integrated polarizer. Samples were excited at 480 nm and emission readings were collected at 535 nm wavelengths. Reactions were carried out at 30 °C in reaction buffer (20 mM Tris-HCl, pH 7.5, 100 mM NaCl and 2 mM DTT) with a total reaction volume of 100 μl. Each reaction mixture contained 50 nM Fluorescein-labeled (6-carboxyfluorescein) DNA substrates with increasing concentrations of Shu complex. Titrated reactions were incubated for 7.5 min before measurements were taken. Change in fluorescence polarization was obtained by subtracting each measurement by that of DNA alone. Triplicate data sets were fit for a one-site binding model, in PRISM5 (GraphPad), to a quadratic equation (**1**) to determine the *K_d_* values.

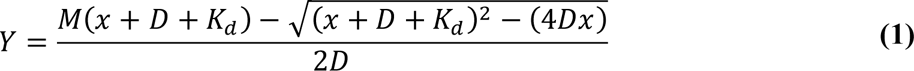

### Protein-ATP binding analysis with a fluorescent ATP Analog

Protocols for the binding assays were adapted from a technique by LaConte *et al*.(38) Experiments were performed using a Synergy H1 Hybrid Multi-Mode Microplate Reader (BioTek Instruments) at 30 °C. Reaction mixtures (100 μl) contained 4 μM protein and 5 μM TNP-ATP in the reaction buffer (50 mM Tris, pH 7.2, 50 mM KCl, 2 mM MgCl_2_ and 2 mM DTT). Buffer alone and buffer with 4 μM lysozyme were used as blank controls. Samples were excited at 410 nm and Fluorescence readings were collected at 540 nm.

For the TNP-ATP saturation assays, emission readings at 540 nm were collected for 2 μM Shu protein samples with increasing TNP-ATP concentrations. The correction factors for the inner filter effect were established and applied to data points above 10 μM TNP-ATP as previously described (39–41). Triplicate data sets were analyzed in PRISM (GraphPad) by non-linear regression with a one-site specific binding model to determine *K_d_^TNP-ATP^*, using the equation (**2**).

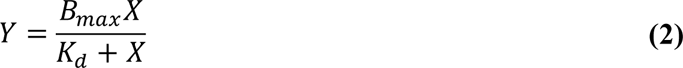

For the competitive binding curves, 2 μM Shu protein with fixed 2.5 μM TNP-ATP was titrated by increasing concentrations of ATP. Triplicate data sets were analyzed in PRISM (GraphPad) by non-linear regression with a one-site competitive binding model to determine the *K_i_^ATP^*, using the equations (**3**) and (**4**).

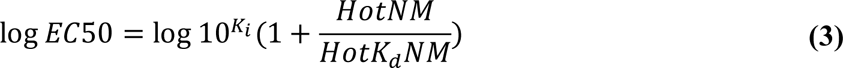

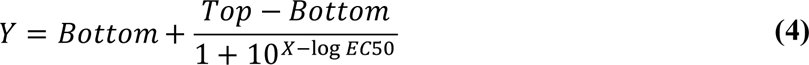

The *K_i_^ATP^*is then used to derive the corresponding *K_d_^ATP^* parameter, using the equations (**5**) and (**6**)(42).

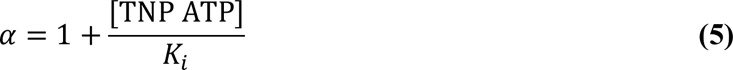

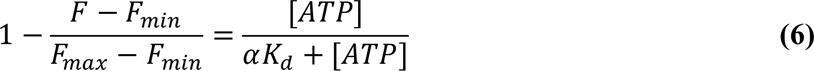

### ATPase Activity Assays

ATPase activity was determined using a malachite green-based assay(43, 44). The assays were performed in 50-μl reaction mixtures containing 50 mM Tris-HCl, pH 7.5, 50 mM NaCl, 5 mM MgCl_2_, 1 mM ATP and 2 mM DTT with 1 μM Shu proteins in the presence and absence of 10 μM DNA at 30 °C. Reactions containing no protein were performed to generate a background reading of inorganic phosphate (Pi) and were subtracted from the experimental results. The inorganic phosphate released was calculated based on the absorbance standard curve established by KH_2_PO_4_ standards.

Steady state ATPase kinetic assays with the Shu complex or Rad51 were performed with/without 1 μM DNA, 1 μM protein and increasing concentrations of ATP in 50-μl volumes. Triplicate data sets were used to calculate the kinetic parameters *K*_m_, *V_max_* and *k*_cat_ with a nonlinear regression fit of the nanomoles Pi released/min to the Michaelis–Menten equation (**7**).

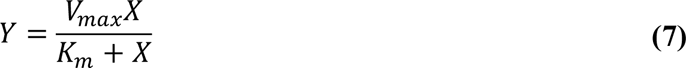

### Nuclease Protection Assays

For standard DNase I protection assays, protein-DNA complexes were formed with different concentrations of Shu complex and 0.25 μM 5’-or 3’-6-FAM-labeled ssDNA substrates in 50 mM Tris, pH 7.5, 50 mM NaCl, 10 mM MgCl_2_, 2 mM ATP (or the indicated nucleotides) and 2 mM DTT. After 30 min at 24 °C, 1 mM CaCl_2_ and the indicated volumes of 2 units bovine pancreatic DNase I (New England Biolabs) were added for the indicated incubation times at 37 °C. For reactions with Rad51, Rad51-ssDNA was incubated for 10 min at 24 °C prior to the addition of the Shu complex, and further incubated for 30 min. For time course experiments, the Shu complex and DNase I, in a mixture with the inclusion of 1 mM CaCl_2_, were added to indicated samples simultaneously. Reaction mixtures (10 μl each) were stopped by the addition of 1 μl of 50 mM EDTA and 12 μl of formamide and resolved in 22.5 % denaturing polyacrylamide urea gels in 1X TBE (300 V, 50 min). All assays were repeated at least three times.

### Fluorescence Assays of Shu-Rad51 Filaments

Experiments were performed using a Synergy H1 Hybrid Multi-Mode Microplate Reader (BioTek Instruments). Samples were excited at 490 nm and fluorescence measurements were collected at the emission wavelength of 522 nm at 30°C. Reactions (100 μl) were carried out in a buffer of 50 mM Tris-HCl, pH 7.5, 5 mM MgCl_2_, 50 mM NaCl and 2 mM DTT. Standard reactions consisted of pre-formed Rad51-ssDNA filaments with 1 μM Rad51 and 15 nM of 5’-or 3’-6-FAM-labeled poly dT39 ssDNA in the presence of 2 mM ATP. The Shu complex (50 nM), pre-incubated with/without the indicated nucleotides, was then incubated together with the pre-formed Rad51-ssDNA filaments to study its influence. In competition assays for simulating filament dissociation, a 100-fold excess of unlabeled poly dT39 ssDNA (1.5 μM) was mixed with the Rad51 filament mixture after the initial 10-min incubation. Buffer controls (with and without DNA) were conducted with each experiment trial to confirm fluorescence signal stability over the time course. Fluorescence readings of labeled ssDNA in buffer were used for background subtraction of all data sets. The initial fluorescence measurement for the pre-formed Rad51-ssDNA filament alone was used as the 0-s time point. Subsequent time-point measurements were used to normalize all data sets.

## Results

### Shu complex’s DNA-binding preferences

The DNA-binding properties of the Shu tetramer complex are important for its role in DNA damage tolerance. Previous studies have reported that the Shu complex and one of its dimers, Csm2-Psy3, bind both ss- and dsDNA with varied binding affinities(27, 31, 32). The Csm2-Psy3 dimer was shown to preferentially bind fork-shaped and 3’-overhang DNA substrates(32). However, there is no systematic analysis for the DNA-binding of the full Shu tetramer. To explore the DNA-binding preferences of the Shu complex, we carried out fluorescence polarization assays (FPAs) with a series of 6-FAM fluorescein-labeled DNA substrates (ss- and dsDNA in different sizes and end types) (**Table S1**). The Shu complex and its mutants in this study were purified to near homogeneity with correct assembling sizes (**Figure 1A & 1B**). We first used Electrophoretic Mobility Shift Assays (EMSAs) to check the physical interactions between WT Shu complex and different DNA substrates. Two-composition native gels were used: a 5 % PAGE layer at the top to allow the large DNA-Shu complex entering the gel and a 15 % PAGE layer at the bottom to prevent the free DNA running out of gels (**Figure 1C**). The EMSA results showed that Shu interacts with both ss- and dsDNA in a concentration-dependent manner (**Figure 1C & S1A**). Then, we carried out fluorescence polarization assays (FPA) to determine the binding affinities of Shu with different DNA substrates (**Figure 1D, 1E & 1F**). The *K_d_* values averaged at ∼3.26 μM for ssDNA, while the *K_d_*values for dsDNA averaged at ∼4.74 μM (**Table 1**). For the ssDNA substrates, *K_d_* values are in a narrow range from 2.37 to 3.54 μM for the ssDNA sizes of 21 to 60 nucleotides (**Figure 1D & Table 1**). For the dsDNA substrates, the *K*_d_ values range from 2.06 to 7.40 μM (**Figure 1E, 1F & Table 1**), much wider than that of ssDNA. The end types, regardless of the lengths of ssDNA component(s), play an important role in binding. Particularly, the blunt-ended dsDNA substrates (ds21, ds26, ds39), regardless of duplex length, were strikingly weaker than those dsDNA with ssDNA component(s) for Shu-DNA binding (**Figure 1E, 1F & Table 1**). Contrary to Csm2-Psy3 dimer’s binding preference for fork-shaped and 3’-overhangs in dsDNA substrates(32), the Shu tetramer favors all the dsDNA with ssDNA components (5’-overhang, 3’-overhang and fork-shaped) (**Figure 1F & Table 1**). There are large disparities in *K_d_* values between ssDNA substrates (ss21, ss26, ss39) and their blunt-ended dsDNA counterparts of the same lengths (ds21, ds26, ds39), with ssDNA substrates having tighter binding affinities than blunt-ended substrates (**Table 1**). Overall, the binding data indicate that the Shu tetramer favors the binding of DNA with ssDNA component(s) over blunt-ended dsDNA. Taken together, our results indicate that the size of DNA substrates is not critical for Shu-DNA binding, but rather the ssDNA components and ssDNA/dsDNA junctions of substrates are the important factors for Shu-DNA interactions.

**Figure 1.**
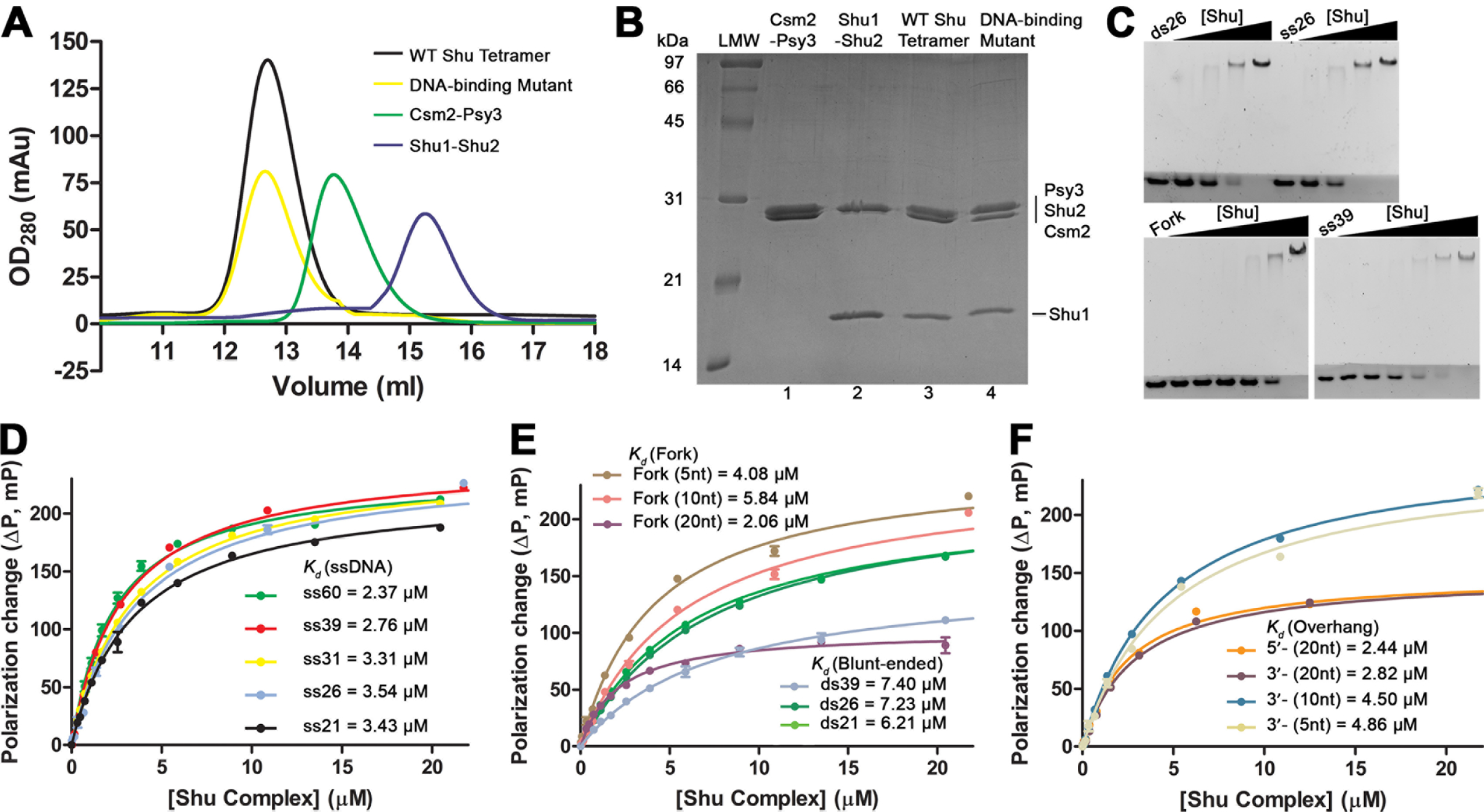
Purified Shu complex and its DNA binding. (**A**) Gel filtration and (**B**) SDS-PAGE analysis of the purified Shu proteins and DNA-binding deficient mutant. (**C**) Electrophoretic Mobility Shift Assay (EMSA) for Shu-DNA interactions. Increasing concentrations of protein (ranging from 0.1 - 3.2 μM) were mixed with 0.2 μM DNA substrates. Protein-DNA mixtures were resolved by 2-layer PAGE gels: 5 % native polyacrylamide at top and a 15 % at the bottom (dark layer). (**D, E, F**) Fluorescence polarization assay (FPA) for DNA-binding of Shu complex with (**D**) ssDNA and (**E, F**) dsDNA substrates. Increasing concentrations of protein were added to 50 nM fluorescein-labeled DNA substrates. Dissociation constants (*K_d_*) and associated standard error of mean from triplicate experiments were determined by non-linear curve fitting to a one-site binding model (**Materials and Methods**). (**E, F**) Number in parentheses of substrate names stands for the number of nucleotides in its single-stranded part(s) from the ssDNA/dsDNA junction.

**Table 1.** DNA-binding affinity of Shu complex.

| DNA <sup>a, b</sup> | Type | Size (nt/bp) <sup>c</sup> | $K_d$ (μM) <sup>d</sup> |
| --- | --- | --- | --- |
| ss21 | ssDNA | 21/- | $3.43 \pm 0.16$ |
| ss26 | ssDNA | 26/- | $3.54 \pm 0.24$ |
| ss31 | ssDNA | 31/- | $3.31 \pm 0.07$ |
| ss39 | ssDNA | 39/- | $2.76 \pm 0.08$ |
| ss60 | ssDNA | 60/- | $2.37 \pm 0.13$ |
| ds21 | Blunt-ended | 21/21 | $6.21 \pm 0.19$ |
| ds26 | Blunt-ended | 26/26 | $7.23 \pm 0.14$ |
| ds39 | Blunt-ended | 39/39 | $7.40 \pm 0.51$ |
| Fork (5nt) | Fork-shaped | 26/21 (5-nt ss) | $4.08 \pm 0.19$ |
| 3'-overhang (5nt) | 3'-overhang | 26/21 (5-nt ss) | $4.86 \pm 0.28$ |
| Fork (10nt) | Fork-shaped | 31/21 (10-nt ss) | $5.84 \pm 0.32$ |
| 3'-overhang (10nt) | 3'-overhang | 31/21 (10-nt ss) | $4.50 \pm 0.12$ |
| Fork (20nt) | Fork-shaped | 39/19 (20-nt ss) | $2.06 \pm 0.13$ |
| 3'-overhang (20nt) | 3'-overhang | 39/19 (20-nt ss) | $2.82 \pm 0.19$ |
| 5'-overhang (20nt) | 5'-overhang | 39/19 (20-nt ss) | $2.44 \pm 0.20$ |

<sup>a</sup> Sequence of DNA substrates can be found in **Table S1**.
<sup>b</sup> Number in parentheses of substrate names stands for the number of nucleotides in its single-stranded part(s) from the ssDNA/dsDNA junction.
<sup>c</sup> “nt” stands for longest DNA-strand size; “bp” stands for duplex size of dsDNA; “ss” stands for ssDNA component of dsDNA substrate.
<sup>d</sup> Data were obtained through non-linear regression of the binding data as described in **Materials and Methods** and represent the mean ± SEM (standard error of mean) of three independent experiments.

We also conducted FPAs on the Csm2-Psy3 and Shu1-Shu2 dimers, and a DNA-binding deficient Shu tetramer mutant with different DNA substrates to compare and validate the DNA binding of Shu. The DNA binding mutant have the positive residues mutated along the putative DNA-binding cleft (K199A/R200A/R201A on Psy3; K189A/R190A/R191A/R192A on Csm2), based on the structure of the Psy3-Csm2 dimer(27). The FPA data show that the Csm2-Psy3 dimer binds ss- and dsDNA with similar affinities to that of Shu (**Figure S2 & Table S2**) while Shu1-Shu2 dimer and DNA-binding mutant do not bind DNA (**Figure S1C)**, which is consistent with the previously study (27). The *K_d_* values for ssDNA and blunt-ended dsDNA substrates are comparable to those reported by Tao *et al*(27). In addition, we determined the DNA-binding affinity of Shu in the presence of ATP, ADP and AMP-PNP, and found that the type of nucleotides does not affect DNA binding of Shu proteins significantly (**Figure S2 & Tables S3 & S4**).

Overall, the binding data served the purpose of systematic comparisons of Shu binding with different types of DNA substrates. Interestingly, Shu shares the similar binding preference of ssDNA over blunt-ended dsDNA (*K*_d_: 3.03 μM vs. 8.54 μM) with Rad51 (*K*_d_: 0.28 μM vs. 2.79 μM) even though Shu binds DNA much weaker than Rad51(45, 46). Noticeably, Rad51 binds ssDNA much tighter than Shu, which makes the binding-affinity disparity between ssDNA and dsDNA ∼10-fold; instead of ∼3-fold in Shu.

### Shu complex binds ATP

Structural analyses revealed that the Psy3 subunit of the Shu complex contains an ATP hydrolysis domain (Walker B motif)(26, 27). RAD51 homologs have the conserved Walker A and Walker B motifs for ATP binding and ATP hydrolysis, respectively(47–50). Through sequence analysis, we found a putative ATP-binding motif (Walker A motif) in Psy3 (**Figure 2**). We examined the ATP-binding of the Shu proteins first, using a fluorescent ATP analog, 5 μM 2’(3’)-O-(2,4,6-trinitrophenyl) adenosine 5’-triphosphate (TNP-ATP). We measured the TNP-ATP fluorescence intensity changes by adding the Shu proteins (the Shu tetramer, Csm2-Psy3 dimer, and Shu1-Shu2 dimer) individually. Both the Shu tetramer and Csm2-Psy3 dimer increased the fluorescence intensity with blue shifts of the emission peaks from 561 nm to 543 nm and 548 nm, respectively (**Figure 3A**). Noticeably, the fluorescence spectra of the Shu tetramer almost superimposable with that of Csm2-Psy3 with a slightly greater blue shift than the Csm2-Psy3 dimer (**Figure 3A**). Nevertheless, the observations indicated that the Shu tetramer and Csm2-Psy3 dimer both bind TNP-ATP and sequester it from aqueous solution. In contrast, the Shu1-Shu2 dimer did not cause any fluorescence changes and essentially overlapped with the buffer baseline’s data set (**Figure 3A**). This suggests that Shu1-Shu2 does not have ATP-binding capabilities. Taken together, the similar fluorescence spectra between Shu tetramer and Csm2-Psy3 suggest that Csm2-Pys3 is the main contributor to the ATP binding in Shu. To validate the ATP binding, we carried out site-specific mutagenesis to mutate the conserved Lys 52 residue in the putative Walker A site (**Figure 2B**). The K52A mutant’s binding spectrum overlapped with that of the buffer control (**Figure 3A**), indicating that the mutation abolished the nucleotide binding of Shu. Thus, the mutagenesis work indicated that the Walker A motif in the Psy3 subunit is responsible for ATP binding in the Shu complex. The proteins in use were well-purified and properly folded with correct elusion positions from size exclusion columns (**Figure 1A, 1B & S1D)**.

**Figure 2.**
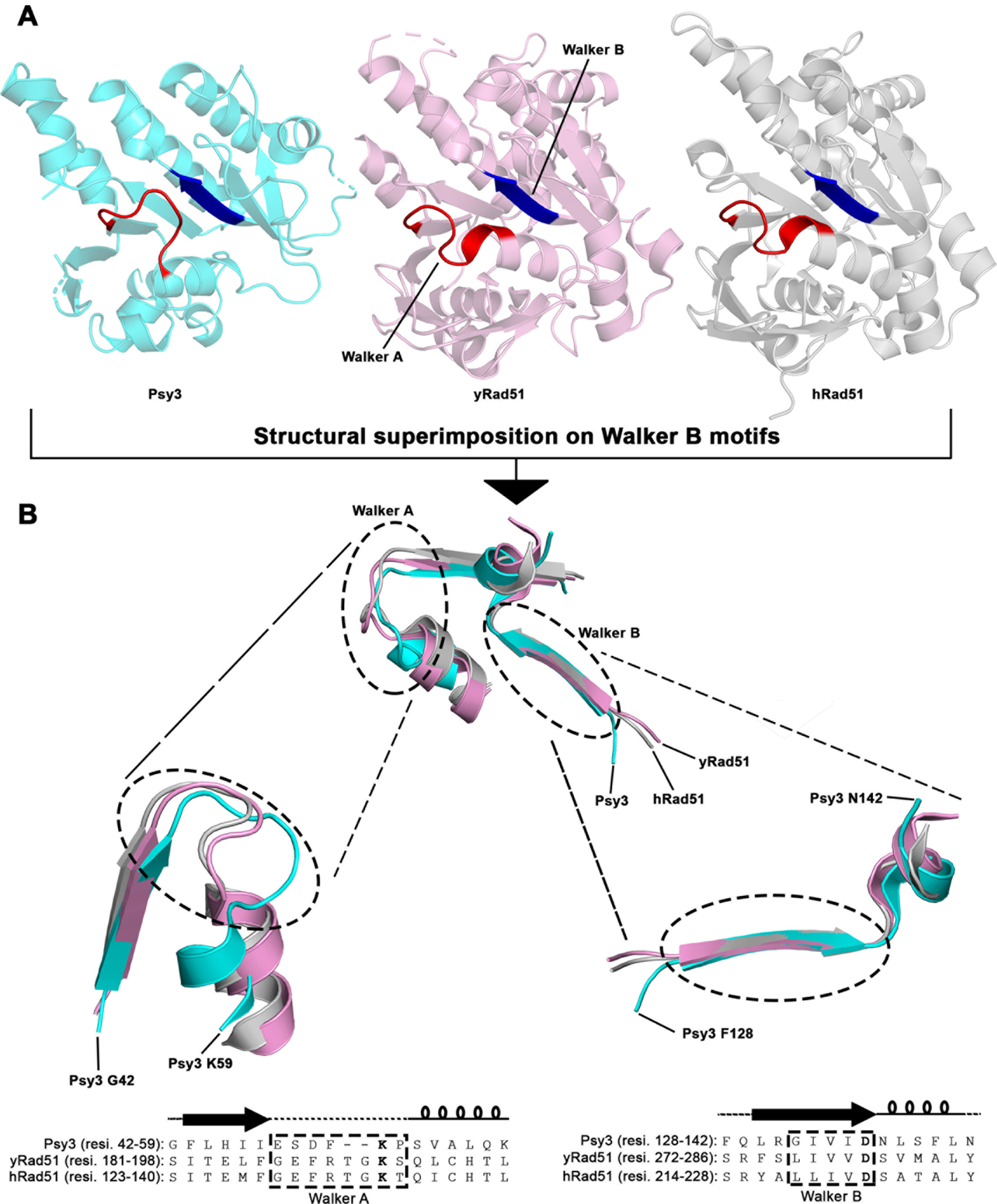
Conserved ATPase Walker A and Walker B motifs in Psy3 of Shu. (**A**) Structural similarity of Psy3 (*cyan*; PDB 4EQ6), yeast Rad51 (*pink*; PDB 1SZP) and human Rad51 (*grey*; PDB 5NWL); Walker A motifs are in *red*. Walker B motifs are in *blue*. (**B**) Structure and sequence alignment of Walker A and B motifs of Psy3, Yeast Rad51 and human Rad51, in same color scheme as in (**A**). The invariant residues for the Walker motifs are bolded in the sequence alignments. Secondary structures of Psy3 are indicated above the sequence alignments with the following: coils, arrows and straight lines represent alpha-helices, beta sheets and loops, respectively.

**Figure 3.**
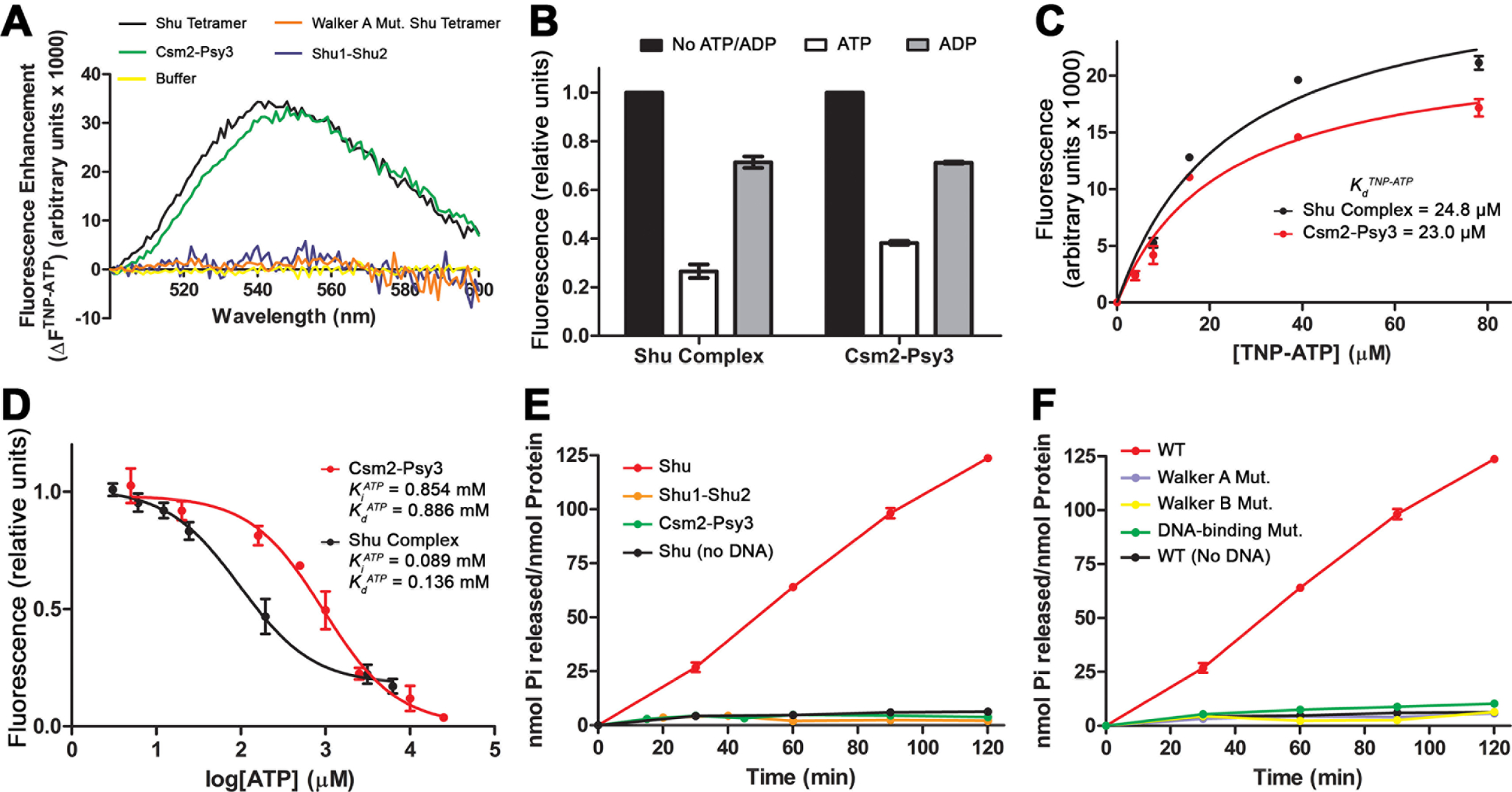
ATP Binding and ATPase activity of Shu. (**A**) Fluorescence spectra of TNP-ATP (5 μM) with the Shu proteins (4 μM). (**B**) Competitions of TNP-ATP (5 μM)/ Shu proteins (4 μM) complexes by ATP or ADP (10 mM each). Fluorescence units were normalized to the initial readings without competitive ATP and ADP. (**C**) TNP-ATP-binding with Shu complex or Csm2-Psy3 in Saturation Assay. Increasing concentrations of TNP-ATP were added to the indicated Shu proteins (2 μM each). The TNP-ATP fluorescence of the samples is plotted after subtracting background TNP-ATP fluorescence obtained with control samples containing lysozyme instead of Shu proteins. Dissociation constants (*K_d_^TNP-ATP^*) were determined by non-linear curve fitting to a one-site binding model. (**D**) Titration of TNP-ATP (2.5 μM) fluorescence with ATP in the presence of Shu or Csm2-Psy3 (2 μM each). Data shown represents fluorescence units after subtraction of the TNP-ATP background fluorescence followed by normalization to the reading in the absence of ATP (relative units). Inhibitory constants (*K_i_^ATP^*) were determined by non-linear curve fitting to a one-site competitive binding model. Dissociation constants (*K_d_^ATP^*) were derived as described in Materials and Methods. (**E**) ATPase activity of the Shu complex and its dimer proteins (1 μM each) in the presence of Fork-shaped DNA substrate (10 μM) over 120-min time course, the tetramer is also tested without DNA. (**F**) ATPase activity of the Shu mutants (1 μM each) with the same DNA substrate in (**E**). The WT Shu data were plotted as a reference.

We further characterized the nucleotide-binding of the Shu complex using ATP and ADP to compete for the binding of TNP-ATP. Adding ATP or ADP (10 mM) in excess to TNP-ATP (5 μM) reduced fluorescence intensities in both the Csm2-Psy3 dimer and Shu tetramer (**Figure 3B**). Noticeably, the fluorescence reductions caused by ATP were greater than ADP, showing a binding preference for ATP. Next, we determined the binding constants of TNP-ATP (*K*_d_^TNP*-*ATP^) for the Csm2-Psy3 dimer (23.0 ± 4.3 μM) and Shu complex (24.8 ± 4.3 μM), respectively (**Figure 3C**); and the binding constants of ATP (*K*_d_^ATP^) for the dimer (0.886 ± 0.290 mM) and tetramer (0.136 ± 0.047 mM), respectively (**Figure 3D**). Interestingly, the Shu complex bound ATP ∼6-fold tighter than the Csm2-Psy3 dimer. This observation indicates that formation of the Shu tetramer contributes to the ATP binding even though Shu1-Shu2 does not bind to ATP directly. Still, the nucleotide-binding affinity of Shu is relatively weak compared to the *K_d_^ATP^* (8.8 μM) of yeast Rad51(51). This suggests that, in a system involving both proteins, Rad51 filaments would likely be saturated with ATP before the Shu complex. Nevertheless, the Shu complex specifically binds ATP/ADP through the Csm2-Psy3 dimer with a Walker A-like motif in the Psy3 subunit.

### Shu complex is a DNA-dependent ATPase

With confirmation ATP binding of the Shu complex, we then explored if it also has ATPase activity. Using malachite green-based ATP hydrolysis assays, we tested the ATPase activity of the Shu proteins. The assays showed that the Shu tetramer complex was able to hydrolyse ATP, while the individual heterodimers (either Shu1-Shu2 or Csm2-Psy3) do have any ATPase activities (**Figure 3E**). Furthermore, the ATPase activity of Shu is dependent on the presence of DNA (**Figure 3E**). To validate this observation, we tested the ATPase activity of the DNA binding mutant of Shu. Expectedly, the DNA-binding mutant lost its ATPase activity (**Figure 3F**), as low as Shu in the absence of DNA (**Figure 3E**). Importantly, this mutant’s nucleotide binding was undisturbed (**Figure S1D**). The results indicate that DNA is a crucial component for the ATPase activity of the Shu complex, which is common for Rad51 paralogs (51, 52).

To prove the ATPase activity of Shu, we made a site-specific mutation on the catalytic Walker B domain of Psy3. We generated a Walker B mutant with a D136A mutation in Psy3, as D136 is the key catalytic residue conserved in Walker B domains based on sequence and structure alignment (**Figure 2B**)(26, 27, 30, 53). Both Walker A and Walker B mutants were purified and examined by sizing column for correct assembly (**Figure S1E**). The ATPase assays on the Walker B mutant were carried out in parallel with the Walker A mutant (ATP binding mutant), DNA-binding mutant and WT Shu. The D136A mutant lost ATPase activity completely (**Figure 3F**), indicating that the activity is specifically from the Walker B domain of Psy3 and intrinsic in Shu. The Walker B mutant maintained its DNA-binding (**Figure S1A & S1F**) and nucleotide-binding abilities (**Figure S1D**). Overall, all three mutants (Walker A, Walker B and DNA-binding mutants) lost their ATPase activities (**Figure 3F**). The results indicate that Shu possesses DNA-dependent ATPase activity, which is an important property for Rad51 paralogs(51, 52).

To quantitatively measure Shu’s ATPase activity, we determined the kinetic parameters of ATP hydrolysis in the presence of different DNA. The *k*_cat_ were in the range of 1.29 - 2.76 min^-1^, representing a slow ATPase which likely plays regulatory roles (**Table 2 & Figure S3A**). In addition, Shu’s ATPase activity is also influenced by the type of DNA. The 5’-overhang and fork-shaped dsDNA substrates had slightly higher *k*_cat_ values than those of the 3’-overhang dsDNA and the ssDNA substrates followed by the blunt-ended dsDNA substrate (**Table 2**). The *k*_cat_ values are correlated with the DNA-binding preference of Shu (**Figure 1D-F & Table 1**); and indicate the importance of a 5’-overhang to the ATP hydrolysis of Shu. The catalytic efficiencies (*k*_cat_/*K*_m_) of different DNA substrates also showed about 2-fold difference between ssDNA/dsDNA with a ssDNA component (range of 32.4 - 44.4) and blunt ended dsDNA (21.4) (**Table 2**), signifying the importance of the ssDNA component for the ATPase activity of Shu.

**Table 2.** ATPase activity of Shu complex.

| DNA <sup>a, b</sup> | Type | $K_m$ (mM) <sup>c</sup> | $V_{max}$<br>($\mu\text{M Pi} \cdot \text{min}^{-1}$) <sup>c</sup> | $k_{cat}$ ($\text{min}^{-1}$) <sup>d</sup> | $k_{cat}/K_m$<br>($\text{s}^{-1} \cdot \text{M}^{-1}$) |
| --- | --- | --- | --- | --- | --- |
| - | - | $4.20 \pm 0.55$ | $0.17 \pm 0.01$ | 0.17 | 0.7 |
| ss39 | ssDNA | $0.67 \pm 0.13$ | $1.65 \pm 0.12$ | 1.65 | 41.2 |
| ds39 | Blunt-ended | $1.00 \pm 0.26$ | $1.29 \pm 0.15$ | 1.29 | 21.4 |
| Fork (20nt) | Fork-shaped | $1.12 \pm 0.28$ | $2.75 \pm 0.32$ | 2.75 | 41.0 |
| 3'-overhang (20nt) | 3'-overhang | $0.72 \pm 0.15$ | $1.92 \pm 0.15$ | 1.92 | 44.4 |
| 5'-overhang (20nt) | 5'-overhang | $1.42 \pm 0.21$ | $2.76 \pm 0.21$ | 2.76 | 32.4 |

<sup>a</sup> Sequence of DNA substrates can be found in **Table S1**.
<sup>b</sup> Number in parentheses of substrate names stands for the number of nucleotides in its single-stranded part(s) from the ssDNA/dsDNA junction.
<sup>c</sup> Data were obtained through non-linear regression of the ATPase data as described in **Materials and Methods** and represent the mean ± SEM (standard error of mean) of three independent experiments.
<sup>d</sup> $k_{cat} = V_{max}/[\text{Shu}]$ , where $[\text{Shu}] = 1 \mu\text{M}$ .

The Shu complex hydrolyse ATP faster (*k_cat_*: 1.29-2.76 min^-1^, **Table 2**) than the yeast Rad51 (yRad51), which has *k_cat_* values in the range of 0.05-0.7 min^-1^ in literature (52, 54). To exclude the deviations caused by experimental conditions, we carried out ATPase activity assays for purified yRad51 in parallel with our Shu analysis. We determined the yRad51 *k_cat_* values in the range of 0.07-0.50 min^-1^ with different DNA substrates (**Table S5, Figure S3B**), which are in the range of data previously reported(54–56). Thus, the Shu complex is a faster ATPase than yRad51 and would dominate ATP hydrolysis in the consortium action of Shu and Rad51 in cells.

### Shu alters ssDNA sensitivity to nuclease digestion

As we have shown that ssDNA is an essential component of Shu-DNA binding, we used DNase I digestion protection assays to probe the changes of ssDNA bound by the Shu complex. DNase I is a nuclease that non-specifically cleaves DNA. However, its activity with ssDNA is much reduced compared to dsDNA (∼500-fold reduction), and it has some DNA sequence preferences(57, 58). To avoid potential sequence biases, we used poly dT (30nt or 39 nt) ssDNA substrate (dT30, dT39), labeled with 5’-6-FAM, for the digestion experiments. We examined the effect of varying the concentration of the Shu complex on DNase I digestion of ssDNA in the presence of ATP at different time points. Unexpectedly, instead of protecting ssDNA, the Shu complex promoted DNA digestion (**Figure 4A**). After a two-hour digestion, DNase I alone did not cleave the poly dT substrates (**lane 2 in Figure 4A**). When the Shu complex was added into the digestion mixture, the substrates began to be digested over time (0.5 to 2 hours), and the levels of digestion correlated with the Shu complex concentrations (**Figure 4A**). The higher the Shu complex concentration, the greater the digestion of the poly dT ssDNA substrate by DNase I. This experiment revealed that the Shu complex could facilitate DNase I cleavage of ssDNA, which resulted in the sensitization of the DNA substrate to DNase I. The concentration dependency of the Shu complex indicates that Shu sensitizes the DNA to DNase I digestion specifically. As we observed ssDNA binding to the Shu complex, the protein-DNA interaction would be responsible for the DNase I sensitization. The results suggest that the specific Shu-DNA interaction either changes the ssDNA conformation to allow greater ease of access or provides structural support to stabilize the scissile site for efficient DNase I cleavage. To validate the observations, we used three Shu mutants (Walker A, Walker B and DNA bindingdeficient mutants) in the DNase I digestion on dT30 ssDNA. Over the two-hour digestion, DNase I digested the DNA substrate substantially in the presence of WT Shu (**lane 3 Figure 4B**). In contrast, all three mutants failed to sensitize the digestion (**lanes 4, 5 & 6 Figure 4B**). The mutant data indicate that the DNA manipulation activity is specific from Shu. Furthermore, the results suggest that Shu’s action on ssDNA is ATP hydrolysis dependent as all three mutants are devoid of ATPase activity (**Figure 3F**).

**Figure 4.**
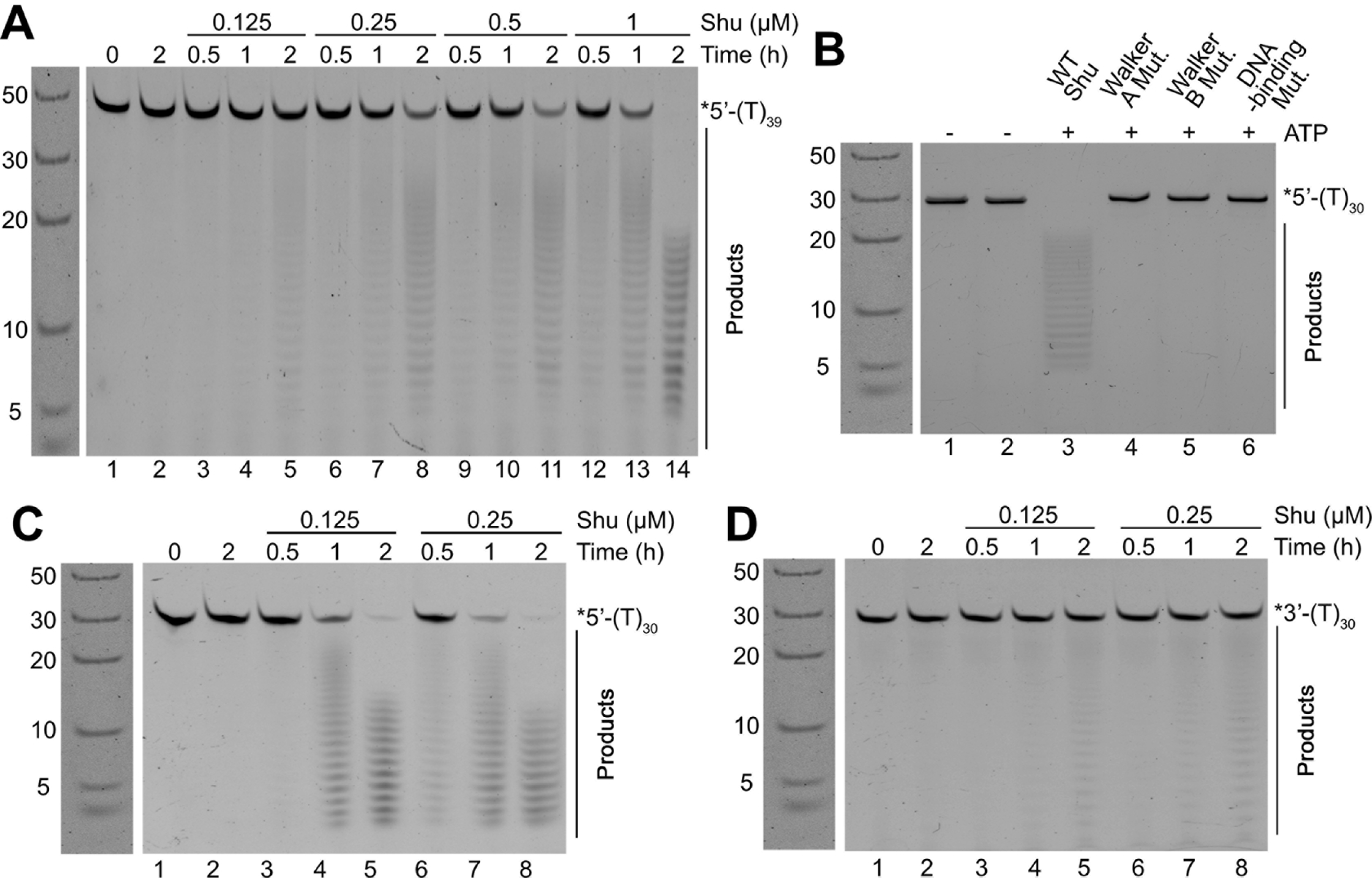
Shu complex enhances DNase I digestion on ssDNA with a 5’ end preference. (**A**) DNase I digestion on 5’-labeled poly dT39 ssDNA (0.25 μM) incubated with Shu complex (0.125 – 1.0 μM), and ATP (2 mM). (**B**) Validation of DNase I digestion enhancement in (**A**) using Shu mutants (1 μM each) and ATP (2 mM) for two hours. (**C, D**) DNase I digestion on (**C**) 5’-or (**D**) 3’-labeled poly dT30 ssDNA (0.25 μM) mixed with Shu complex (0.125 – 0.25 μM) and ATP. Digestions were carried out using 0.5 Unit DNase I at 37 °C. Reaction mixtures were resolved by 22.5 % denaturing PAGE.

To explore the polarity of the action from Shu, we analysed DNase I sensitization of poly dT30 ssDNA substrates labeled at the 5’ and 3’ ends, respectively. Degradation of ssDNA was enhanced for both 5’-and 3’-labeled substrates in the presence of the Shu complex, but at very different rates (**Figures 4C & 4D**). In the presence of Shu, the degradation products of the 5’-labeled substrate were clearly present at the 1-hour time point (**lanes 4 & 7, Figure 4D**) and the full-length substrate completely disappeared after two hours (**lane 5 & 8, Figure 4D**). In contrast, the 3’-labeled digestion products were barely visible over the time (**Figure 4C**). The observations indicate that a preference for the 5’ end in Shu’s action on DNA.

### Shu complex alters accessibility of Rad51-ssDNA filaments

To investigate if the Shu complex acts on Rad51-ssDNA filaments, we carried out fluorescence-based assays on Rad51-ssDNA filaments with labeled ssDNA. We incubated Rad51 with fluorescently labelled poly dT ssDNA (dT39) in the presence of 2 mM ATP for 10 minutes for RAD51-ssDNA filament formation. Then, we measured fluorescence intensities of the Rad51 filaments with/without additions of Shu at 10-second intervals for a 60-second time frame. This assay is used to check the initial effects of Shu on Rad51 filaments(18, 33). Without Shu, the Rad51 filaments have stable fluorescence intensities which were normalized as a baseline (dashed black, **Figure 5A**). The fluorescence intensities of the 5’-labeled Rad51-ssDNA filaments were over 50 % reduced within 10 seconds after Shu complex addition (**red line in Figure 5A**), while no reduction was observed in the 3’-labeled sample within the 60-second time frame (**green line in Figure 5A**). The fluorescence signals of the filaments changed within the first 10 seconds and remained stable from the 10-second point to the 60-second point (**Figure 5A**). The observations reveal that the alteration occurs only at the 5’ end within the first 10-second, but no changes at the 3’ end of the filament up to 60 seconds. The preference of Shu on the 5’ end of the filaments is consistent with its 5’ preference for ssDNA discussed above. The alteration of Rad51 filaments could be caused by either weakened ssDNA binding or conformational changes of the filaments. However, it is unlikely that the Shu complex disrupts the ssDNA binding of Rad51, as Rad51 binds ssDNA (*K_d_* ≈ 0.28 μM)(45) 10-fold tighter than the Shu complex (*K_d_*≈ 2.8 μM). Thus, the cause for the fluorescence reduction is likely conformational changes of the filaments, which makes the fluorophore more solvent exposed than that of the original Rad51 filaments. The polarity indicates that the Shu complex acts at the 5’ end first and alters Rad51 filament conformation at the 5’ end of Shu-Rad51-ssDNA filament. Taken together, the Shu complex binds on Rad51 filaments at the 5’ end initially. To validate the observation above, we used three Shu mutants (DNA binding, Walker A and Walker B mutants) to repeat the fluorescence assays as negative controls. Indeed, the fluorescence intensities of the filaments by the mutants were not changed and overlapped with the baseline over the 60-second time frame (**Figure 5B**). The Walker motif mutants still possess DNA binding (**Figure S1A & S1F**), while the DNA-binding and the Walker B mutants retained ATP binding (**Figure S1D**). Thus, the loss of ATPase activity in all three mutants (**Figure 3F**) indicates that the ATPase activity of Shu is essential for its action on the Rad51 filaments.

**Figure 5.**
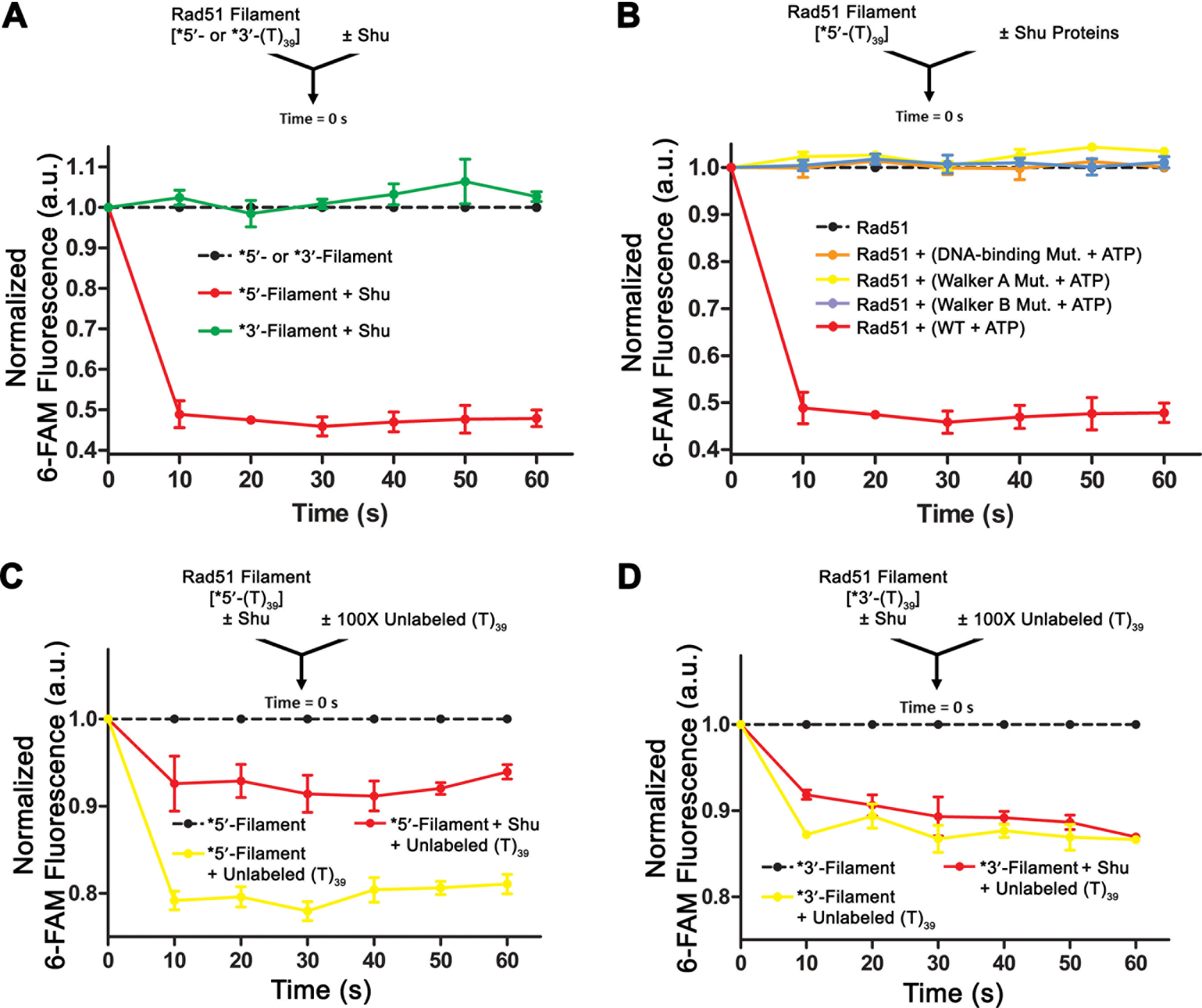
Conformational and stability changes of Rad51 filaments induced by Shu. (**A**) Fluorescence profiles of Rad51 filaments with/without Shu complex over 60-second time. Filaments were pre-formed with Rad51 (1 μM) incubated with 5’-or 3’-labeled poly dT39 ssDNA (15 nM each), then mixed with Shu complex proteins (50 nM each), with ATP (2 mM). (**B**) Fluorescence profiles of 5’-labeled ssDNA-Rad51 filaments with Shu mutants. The WT Shu data in (**A**) were plotted as a reference. (**C, D**) Fluorescence profiles of Rad51-ssDNA filaments with/without Shu complex, mixed with 100-fold excess unlabeled ssDNA, over 60-second time. The arrow indicates the components that are mixed at 0 s time point. Samples were excited at 490 nm and emission readings were measured at 522 nm at equilibrium (30 °C). Filament alone data were normalized as the baseline in arbitrary units corresponding to fluorescence intensities (dashed black lines).

Next, we analyzed the influence of Shu on Rad51 filament stability. Using pre-formed Rad51 filaments with a fluorescently labeled ssDNA, we measured the fluorescence changes of the Rad51 filaments when mixed with unlabeled ssDNA counterpart. The fluorescence intensities were reduced in 10 seconds and remained at those reduced levels in the testing time frame (60 seconds) when an excess of unlabeled ssDNA was added (**yellow lines in Figures 5C & 5D**). This reflects replacement of labeled ssDNA by the unlabeled ssDNA in the Rad51-ssDNA filament. The reduction levels were varied in terms of the absence/presence of Shu and the locations (5’ or 3’ ends) of the fluorescence labels. In the absence of Shu, the fluorescence reduction of 5’-labeled Rad51 filaments was greatest amongst all conditions, indicating the 5’ end of Rad51 filaments is most active for Rad51 filament assembly and dissociation (**yellow line in Figures 5C)**. In the presence of the Shu complex, the fluorescence reduction by the excess unlabeled DNA was partially alleviated for the 5’-labeled substate (**red line in Figure 5C)**, indicating the Shu complex reduced the DNA exchanges. In contrast, the fluorescence reduction for the 3’-labeled filaments remained at similar levels with/without the Shu complex, suggesting that the Shu complex does not significantly affect the 3’ end at this time frame of 60 seconds (**Figure 5D**). Overall, the observations indicate that the Rad51 filaments are more dynamic at the 5’ ends than the 3’ ends of the filaments within the tested time frame (60 seconds) and that this dynamic property can be dampened by the Shu complex.

To further characterize the Shu complex’s influences on Rad51-ssDNA filaments, we conducted DNase I digestion assays on Rad51 filaments in the presence/absence of the Shu complex. The results showed that the Shu complex enhanced the digestion for both 5’- and 3’-labeled Rad51 filaments (**lanes 5-8 in Figure 6A & 6B**), which contrasts with the digestion of Rad51 filaments alone (**lane 4 in Figure 6A & 6B**). With increasing concentrations of the Shu complex, the Rad51-ssDNA filaments showed increased sensitivity to DNase I (**lanes 5-8 in Figures 6A & 6B**). These observations indicate that the Shu complex increased the accessibility of the ssDNA along the Rad51 filament specifically, which is consistent with our observation of enhanced solvent exposure of ssDNA in the fluorescence intensities for the Rad51 filaments (**Figure 5A**). In terms of polarity of the sensitization, the 5’-labeled filament was more sensitive to DNase I cleavage than the 3’-labeled one (**Figures 6A & 6B**), indicating the preference of the 5’-end. However, when the concentration of the Shu complex was increased to reach a DNA-to-Shu molar ratio of 1:8 with Rad51 filament (also in 8-fold excess to DNA), the levels of relative sensitization became similar for both 5’- and 3’-labeled substrates (**lanes 8 in Figures 6A & 6B**). The molar ratio effect indicates that the Shu complex can act on the entire filament, but with preference for the 5’ end. The negative control digestion with three mutants indicate the enhancement of digestion of Rad51 filament is indeed from the WT Shu complex that is capable of hydrolyzing ATP (**Figure 6C**). Overall, the data suggest that the Shu complex causes changes in Rad51-ssDNA filaments, with preference for the 5’ end over the 3’ end, enhancing the accessibility of the Rad51-coated ssDNA to the DNase I digestion. Again, the Shu complex has a similar 5’ end-preference as in its manipulation of ssDNA. Additionally, the 5’ ends of Rad51 filaments may innately be active and dynamic in nature. Thus, both factors could contribute to the Shu complex’s 5’ end preference when acting on Rad51 filaments.

**Figure 6.**
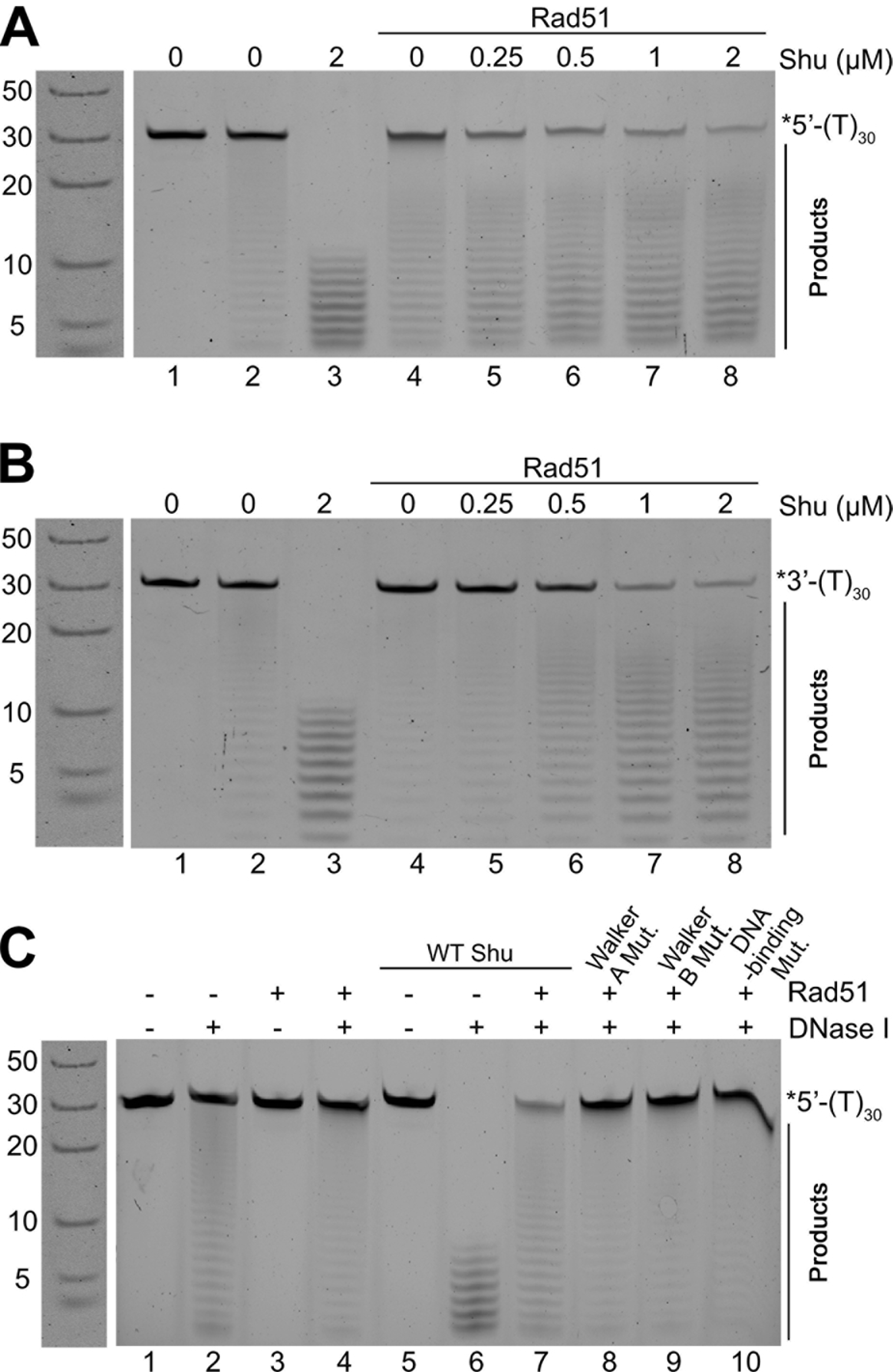
DNase I digestion on Rad51-ssDNA with Shu complex. (**A, B**) Two-hour DNase I digestion on Rad51-ssDNA filaments labeled on (**A**) 5’-or (**B**) 3’-end with increasing concentrations of Shu complex and 2 mM ATP. (**C**) Two-hour DNase I digestion on Rad51 filaments with WT and Shu mutants. Filaments were pre-formed by incubating Rad51 (2 μM) with poly dT30 ssDNA (0.25 μM) and ATP (2 mM) for 10 minutes, then mixed with Shu complex variants (2 μM each). Digestions were carried out using 2 Unit DNase I at 37 °C. Reactions were resolved by 22.5 % denaturing PAGE.

## Discussion

Biochemical characterization of the Shu complex is important to understand its molecular role in DNA damage tolerance. Our quantitative analysis (*K_d_*) on the DNA binding of the Shu complex (tetramer) shows that it preferentially binds DNA substrates with single-stranded component(s), particularly those with a ssDNA/dsDNA junction (characteristic of DNA in HR). The preference of binding ssDNA is similar to that of Rad51 proteins(59, 60). The binding affinities are comparable to the previous work on the Csm2-Psy3 dimer(27, 32), as the dimer is mainly responsible for DNA binding in the Shu complex. However, our Shu tetramer study does not have the DNA-binding preference for 3’-overhangs as does the Csm2-Psy3 dimer(32). Instead, the forked and the 5’-overhang structures of DNA is slightly more favored than the 3*’*-overhang in our binding assays for the Shu tetramer. The Shu complex FPA data showed a clear *K_d_*disparity between the blunt-ended dsDNA substrates and substrates with ssDNA element(s), which is not present in the Csm2-Psy3 data. Thus, it is likely that the complete Shu complex renders better DNA-binding specificity for HR-related DNA substrates than the Csm2-Psy3 dimer.

Our work demonstrate that the Shu complex is capable of binding ATP, and its nucleotide-binding capacity is conferred by the Csm2-Psy3 dimer. Through mutagenesis, we confirmed that the Walker A motif in Psy3’s is responsible for nucleotide binding. Our work further illustrates that the Shu tetramer is a unique DNA-dependent ATPase and validated its ATPase function through mutagenesis on Walker B motif in Psy3 (D136A). ATPase activity has been reported in the human Shu homolog, hSWS1-SWSAP1(16), and is likely a conserved function in Shu homologs. Surprisingly, neither Csm2-Psy3 nor Shu1-Shu2 dimers possess ATPase activity individually. Although both ATP- and DNA-binding functions are solely located in the Csm2-Psy3 dimer, the Shu1-Shu2 dimer is still indispensable for ATPase activity. The Shu2 subunit is unique in the Shu complex, as it contains a conserved SWIM domain. Disruptions of the SWIM domain negatively impact the DNA damage tolerance response in eukaryotes(20), consistent with its indispensability in the Shu complex’s ATPase activity. Thus, the Shu complex’s DNA-dependent ATP hydrolysis is associated with its role in error-free DNA lesion bypass.

We revealed that enhancement of the Shu complex’s ATPase activity by dsDNA substrates with ssDNA components and ssDNA/dsDNA junctions were greater than blunt-ended dsDNA and ssDNA alone. Particularly, we found the 5’-overhang of dsDNA substrates enhanced the ATPase activity of the Shu complex most effectively. Intriguingly, the ATPase activity assays were sensitive the types of DNA, which demonstrates the preference of Shu for 5’-overhang of dsDNA substrates. This suggests that optimal ATPase activity of the Shu complex likely requires very specific binding with a certain type of DNA. Such DNA-type specificity for binding and enzymatic activity stimulation is a common characteristic of Rad51 modulator proteins, such as human BRCA2, a key protein that promotes human Rad51 filament formation(5, 61–64). Furthermore, the Shu complex favors a 5*’*-overhanging structure, which represents the lagging strand of a replication fork. A recent study by Rosenbaum *et al*. suggested that lesions on the lagging strand would be specifically recognized by the Shu complex for HR bypassing(31). The 5*’*-preferences of ATPase stimulation illustrated here are consistent with lesion recognition at stalled replication forks(31). Our work provides important evidence that ssDNA components, ssDNA/dsDNA junctions and the polarity of the overhang in ssDNA/dsDNA duplexes are important for the Shu complex’s function.

We provide evidence of the Shu complex changing ssDNA sensitivity to DNase I with dependency on its ATP hydrolysis activity. It has been suggested that the preference of DNase I to dsDNA over ssDNA is due to the conformational stability of B-form dsDNA, which fits well in the active site of DNase I(57, 58). It seems that the Shu-ssDNA interaction stabilizes ssDNA, causing it to mimic dsDNA at cutting sites. The increase in DNase I sensitization in the presence of ATP over time is coupled with the Shu complex’s ATPase activity (**Figure 4A**).

Interestingly, the Shu complex has a similar effect on Rad51-ssDNA filaments in our DNase I digestion experiments as it does on ssDNA. This Shu-mediated action on Rad51 filaments also demonstrates a 5’-end preference. Our study indicates that the Shu-ssDNA interaction changes the properties of ssDNA and the properties of Rad51 nucleoprotein filaments; the latter was also observed in the studies on the *C*. *elegans* Shu homolog(18, 33). We found that ATP hydrolysis primarily contribute to Shu’s action on the filaments. The influences of ATP hydrolysis to the action of Shu on the filament, again, suggest that the function of Shu can be regulated by its own ATPase activity (**Figure 7A**). The regulation is induced by ATP hydrolysis, which is essential to Shu’s action on Rad51 filaments (**Figure 7B**). Furthermore, the Shu complex’s preference to bind at the 5’ ends of ssDNA and Rad51 filaments fits its role in lesion bypass on the lagging strand of a replication fork(31). In closing, our work illustrates the complex interplay between DNA binding, nucleotide binding, and the ATPase activity of the Shu complex, which helps to regulate Rad51 filaments in HR associated DNA damage responses (**Figure 7**).

**Figure 7.**
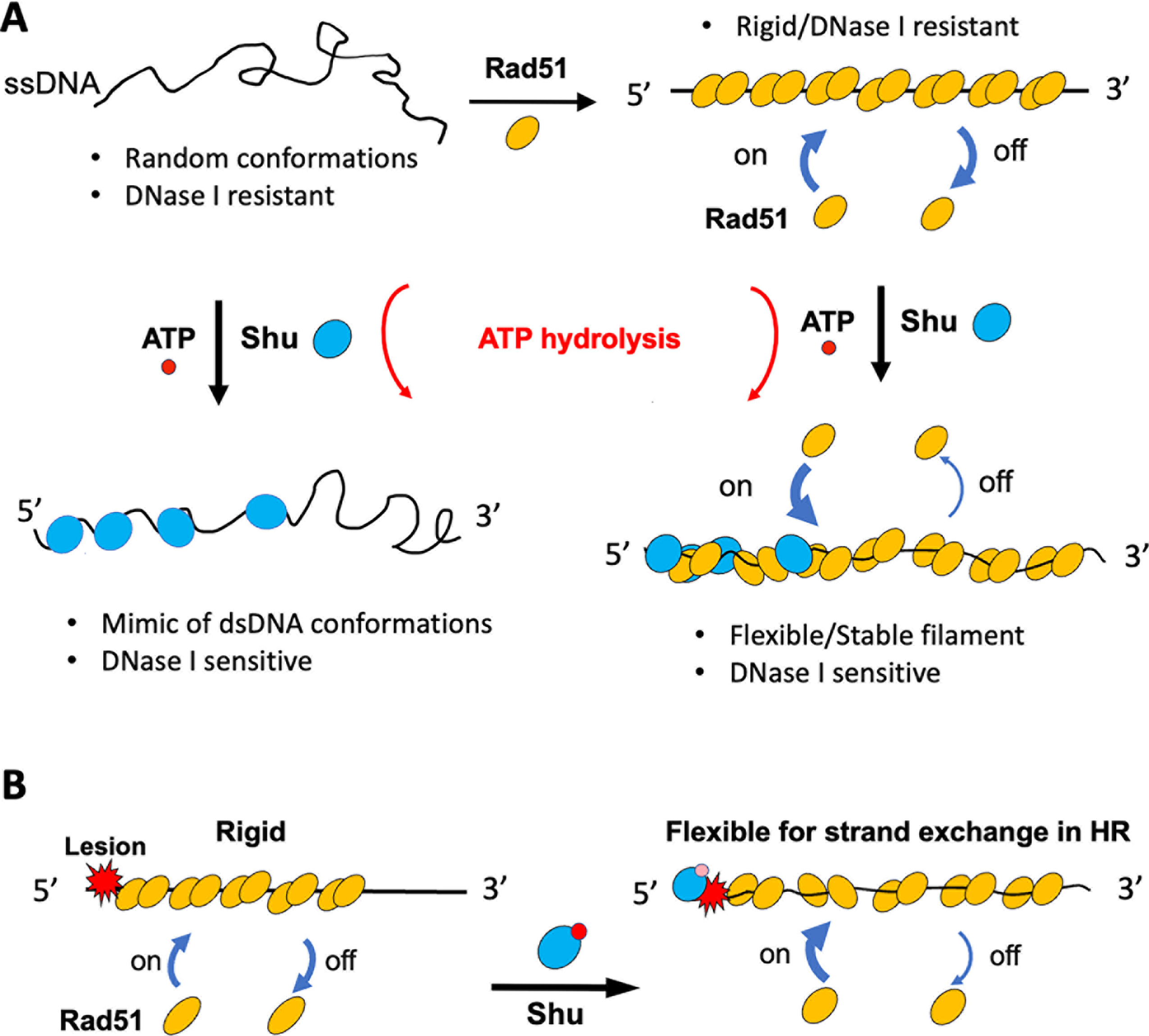
Molecular model for the action of Shu on ssDNA and Rad51 filaments. (**A**) Alteration of Shu on ssDNA and Rad51-ssDNA filaments with 5*’*-to-3*’* polarity; left, Shu binds on ssDNA and makes randomly structured ssDNA in conformations ready for DNase I cleavage; right, Shu binds on Rad51 filaments and changes the conformations of filaments, stabilizing the Rad51 binding on ssDNA and making the filaments flexible/accessible to DNase I. (**B**) The role of Shu in error-free lesion bypass as an DNA-dependent ATPase. Shu binds on the 5*’* end of Rad51 filament which contains lesions, such as abasic lesions, to make the filaments stable and flexible for strand exchange/homologous recombination. The alteration of Rad51 filaments is dependent on the ATPase activity of Shu.

## Data Availability

Source data are available as supplementary files or upon request.

## Acknowledgements

We thank Dr. Wei Xiao for providing original plasmids of the yeast Shu complex, and Dr. Lynn Weir for careful proofreading of the manuscript.

## Abbreviations

Homologous recombination (HR), Shu complex/tetramer (Shu), Methyl Methanesulfonate (MMS), Hydroxyurea (HU), Single-stranded (ss), Double-stranded (ds), Polymerase incomplete primer extension (PIPE), Electrophoretic Mobility Shift Assays (EMSAs), Fluorescence Polarization Assays (FPA), inorganic phosphate (Pi)

