## Supplemental Data for "Shu complex is an ATPase that regulates Rad51 filaments in homologous recombination-directed DNA damage response"

**Supplementary data**

**Table S1. Sequences of DNA substrates.**

| Substrate | Sequence [a] |
|---|---|
| ss21 | 5′- *ACTTACAGCACAGCGGTTTTT -3′ |
| ss26 | 5′- *ACTGCCCTTACAGCACAGCGGTTTTT -3′ |
| ss31 | 5′- *ACTGCCCTTACAGCACAGCGGTTTTTTTTTT -3′ |
| ss39 | 5′- TTTTTTTTTTTTTTTTTTTTCTTGACAAGCTTGCGCACT* -3′ |
| ds21 | 5′- *ACTTACAGCACAGCGGTTTTT -3′<br>3′- TGAATGTCGTGTCGCCAAAAA -5′ |
| ds26 | 5′- *ACTGCCCTTACAGCACAGCGGTTTTT -3′<br>3′- TGACGGGAATGTCGTGTCGCCAAAAA -5′ |
| Fork (5nt) | 5′- *ACTGCCCTTACAGCACAGCGGTTTTT -3′<br>3′- TGACGGGAATGTCGTGTCGCCTTTTT -5′ |
| 3′-overhang (5nt) | 5′- *ACTGCCCTTACAGCACAGCGGTTTTT -3′<br>3′- TGACGGGAATGTCGTGTCGCC -5′ |
| Fork (10nt) | 5′- *ACTGCCCTTACAGCACAGCGGTTTTTTTTTT -3′<br>3′- TGACGGGAATGTCGTGTCGCCTTTTTTTTTT -5′ |
| 3′-overhang (10nt) | 5′- *ACTGCCCTTACAGCACAGCGGTTTTTTTTTT -3′<br>3′- TGACGGGAATGTCGTGTCGCC -5′ |
| Fork (20nt) | 5′- TTTTTTTTTTTTTTTTTTTTCTTGACAAGCTTGCGCACT* -3′<br>3′- TTTTTTTTTTTTTTTTCACGGAACTGTTCGAACGCGTGA -5′ |
| 3′-overhang (20nt) | 5′- CTTGACAAGCTTGCGCACT* -3′<br>3′- TTTTTTTTTTTTTTTTCACGGAACTGTTCGAACGCGTGA -5′ |
| 5′-overhang (20nt) | 5′- TTTTTTTTTTTTTTTTTTTTCTTGACAAGCTTGCGCACT* -3′<br>3′- GAACTGTTCGAACGCGTGA -5′ |
| Fork (20nt) | 5′- TTTTTTTTTTTTTTTTTTTTCTTGACAAGCTTGCGCACT -3′<br>3′- TTTTTTTTTTTTTTTTCACGGAACTGTTCGAACGCGTGA -5′ |
| ss39 | 5′- TTTTTTTTTTTTTTTTTTTTCTTGACAAGCTTGCGCACT -3′ |
| ds39 | 5′- TTTTTTTTTTTTTTTTTTTTCTTGACAAGCTTGCGCACT -3′<br>3′- AAAAAAAAAAAAAAAAAAAAGAACTGTTCGAACGCGTGA -5′ |
| 5′-overhang (20nt) | 5′- TTTTTTTTTTTTTTTTTTTTCTTGACAAGCTTGCGCACT -3′<br>3′- GAACTGTTCGAACGCGTGA -5′ |
| 3′-overhang (20nt) | 5′- CTTGACAAGCTTGCGCACT -3′<br>3′- TTTTTTTTTTTTTTTTCACGGAACTGTTCGAACGCGTGA -5′ |
| 5′-labeled poly dT39 (39-mer) | 5′- *TTTTTTTTTTTTTTTTTTTTTTTTTTTTTTTTTTTTTTT -3′ |
| 5′-labeled poly dT (30-mer) | 5′- *TTTTTTTTTTTTTTTTTTTTTTTTTTTTTT -3′ |
| 3′-labeled poly dT30 with 5′-P (30-mer) | 5′- TTTTTTTTTTTTTTTTTTTTTTTTTTTTTT* -3′ |
| 3′-labeled polydT39 with 5′-P (39-mer) | 5′- TTTTTTTTTTTTTTTTTTTTTTTTTTTTTTTTTTTTTTT* -3′ |
| ss60 | 5′- TTTTTTTTTTTTTTTTTTTTTTTTTTTTTT<br>CTTGACAAGCTTGCGCACTTAGGCCTCTAG* -3′ |

[a] Symbol “*” denotes the position at which a single-fluorescein molecule of 6-FAM is attached.

**Table S2. DNA-binding affinity of Csm2-Psy3 dimer (see Fig. S1B).**

| **DNA** [a, b] | **Type** | **Size (nt/bp)** [c] | **$K_d$ (μM)** [d] |
|---|---|---|---|
| ss39 | ssDNA | 39/- | 1.65 ± 0.18 |
| ds39 | Blunt-ended | 39/39 | 1.49 ± 0.13 |
| Fork (20nt) | Fork-shaped | 39/19 (20-nt ss) | 1.35 ± 0.06 |
| 3′-overhang (20nt) | 3′-overhang | 39/19 (20-nt ss) | 1.92 ± 0.28 |
| 5′-overhang (20nt) | 5′-overhang | 39/19 (20-nt ss) | 1.18 ± 0.10 |

[a] Sequence of DNA substrates can be found in **Table S1**.

[b] Number in parentheses of substrate names stands for the number of nucleotides in its single-stranded part(s) from the ssDNA/dsDNA junction.

[c] "nt" stands for longest DNA-strand size; "bp" stands for duplex size of dsDNA; "ss" stands for ssDNA component of dsDNA substrate.

[d] Data were obtained through non-linear regression of the binding data as described in **Materials and Methods** and represent the mean ± SEM (standard error of mean) of three independent experiments.

**Table S3. DNA-binding affinity of Shu in the presence of nucleotides**

| DNA [a, b] | Type | Size (nt/bp) [c] | Nucleotide | $K_d$ (mM) [d] |
|---|---|---|---|---|
| ss60 | ssDNA | 60/- | - | 2.37 ± 0.13 |
| | | | ATP | 2.37 ± 0.29 |
| | | | ADP | 2.40 ± 0.23 |
| | | | AMP-PNP | 2.01 ± 0.26 |
| ss39 | ssDNA | 39/- | - | 2.76 ± 0.08 |
| | | | ATP | 2.77 ± 0.25 |
| | | | ADP | 3.19 ± 0.27 |
| | | | AMP-PNP | 2.37 ± 0.22 |
| ds39 | Blunt-ended | 39/39 | - | 7.23 ± 0.14 |
| | | | ATP | 7.41 ± 0.61 |
| | | | ADP | 6.85 ± 0.49 |
| | | | AMP-PNP | 5.80 ± 0.40 |
| 3′-overhang (20nt) | 3′-overhang | 39/19 (20-nt ss) | - | 2.82 ± 0.19 |
| | | | ATP | - |
| | | | ADP | 2.53 ± 0.17 |
| | | | AMP-PNP | 2.71 ± 0.22 |
| Fork (20nt) | Fork-shaped | 39/19 (20-nt ss) | - | 2.06 ± 0.13 |
| | | | ATP | 1.03 ± 0.12 |
| | | | ADP | 0.86 ± 0.07 |
| | | | AMP-PNP | 0.97 ± 0.10 |
| 5′-overhang (20nt) | 5′-overhang | 39/19 (20-nt ss) | - | 2.44 ± 0.20 |
| | | | ATP | 1.15 ± 0.11 |
| | | | ADP | 1.58 ± 0.16 |
| | | | AMP-PNP | 2.42 ± 0.22 |

[a] Sequence of DNA substrates can be found in **Table S1**.

[b] Number in parentheses of substrate names stands for the number of nucleotides in its single-stranded part(s) from the ssDNA/dsDNA junction.

[c] "nt" stands for longest DNA-strand size; "bp" stands for duplex size of dsDNA; "ss" stands for ssDNA component of dsDNA substrate.

[d] Data were obtained through non-linear regression of the binding data as described in **Materials and Methods** and represent the mean ± SEM (standard error of mean) of three independent experiments.

**Table S4. DNA-binding affinity of Csm2-Psy3 dimer in the presence of nucleotides (see Figs. S2A, S2B & S2C).**

| DNA [a, b] | Type | Size (nt/bp) [c] | Nucleotide | $K_d$ (mM) [d] |
|---|---|---|---|---|
| ss39 | ssDNA | 39/- | - | 1.65 ± 0.18 |
| | | | ATP | 1.32 ± 0.12 |
| | | | ADP | 1.49 ± 0.18 |
| | | | AMP-PNP | 1.98 ± 0.20 |
| ds39 | Blunt-ended | 39/39 | - | 1.49 ± 0.13 |
| | | | ATP | 1.45 ± 0.13 |
| | | | ADP | 1.54 ± 0.08 |
| | | | AMP-PNP | 1.46 ± 0.12 |
| 3′-overhang (20nt) | 3′-overhang | 39/19 (20-nt ss) | - | 1.92 ± 0.28 |
| | | | ATP | 1.27 ± 0.12 |
| | | | ADP | 1.60 ± 0.13 |
| | | | AMP-PNP | 0.68 ± 0.05 |
| Fork (20nt) | Fork-shaped | 39/19 (20-nt ss) | - | 1.35 ± 0.06 |
| | | | ATP | 0.75 ± 0.07 |
| | | | ADP | 0.74 ± 0.05 |
| | | | AMP-PNP | 0.78 ± 0.05 |
| 5′-overhang (20nt) | 5′-overhang | 39/19 (20-nt ss) | - | 1.18 ± 0.10 |
| | | | ATP | 0.95 ± 0.12 |
| | | | ADP | 0.43 ± 0.07 |
| | | | AMP-PNP | 0.76 ± 0.09 |

[a] Sequence of DNA substrates can be found in **Table S1**.

[b] Number in parentheses of substrate names stands for the number of nucleotides in its single-stranded part(s) from the ssDNA/dsDNA junction.

[c] "nt" stands for longest DNA-strand size; "bp" stands for duplex size of dsDNA; "ss" stands for ssDNA component of dsDNA substrate.

[d] Data were obtained through non-linear regression of the binding data as described in **Materials and Methods** and represent the mean ± SEM (standard error of mean) of three independent experiments.

**Table S5. Rad51 ATPase activity**

| DNA [a, b] | Type | $K_m$ (mM) [c] | $V_{max}$ ($\mu$M Pi · $min^{-1}$) [c] | $k_{cat}$ ($min^{-1}$) [d] | $k_{cat}/K_m$ ($s^{-1} \cdot M^{-1}$) |
|---|---|---|---|---|---|
| - | - | 0.15 ± 0.03 | 0.072 ± 0.002 | 0.07 | 8.21 |
| ss39 | ssDNA | 1.67 ± 0.33 | 0.318 ± 0.037 | 0.32 | 3.18 |
| ds39 | Blunt-ended | 3.65 ± 1.10 | 0.183 ± 0.040 | 0.18 | 0.84 |
| Fork(20nt) | Fork-shaped | 0.79 ± 0.11 | 0.278 ± 0.016 | 0.28 | 5.83 |
| 3'-overhang(20nt) | 3'-overhang | 2.01 ± 0.48 | 0.271 ± 0.040 | 0.27 | 2.25 |
| 5'-overhang(20nt) | 5'-overhang | 5.78 ± 2.74 | 0.501 ± 0.190 | 0.50 | 1.45 |

[a] Sequence of DNA substrates can be found in **Table S1**.

[b] Number in parentheses of substrate names stands for the number of nucleotides in its single-stranded part(s) from the ssDNA/dsDNA junction.

[c] Data were obtained through non-linear regression of the ATPase data as described in **Materials and Methods** and represent the mean ± SEM (standard error of mean) of three independent experiments.

[d] $k_{cat} = V_{max}$/[Rad51], where [Rad51] = 1 $\mu$M.

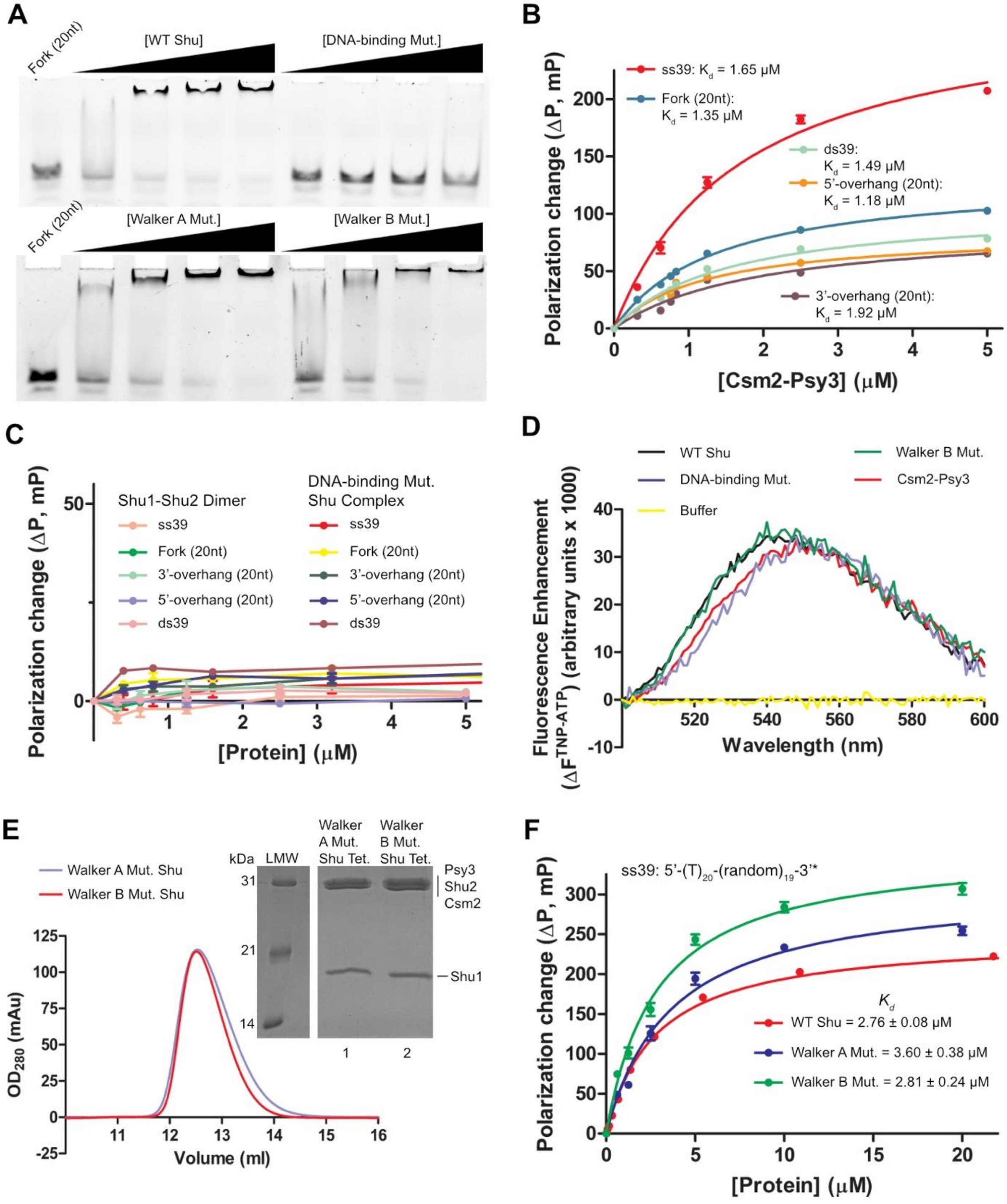
A
Fork (20nt)
[WT Shu]
[DNA-binding Mut.]
[Walker A Mut.]
[Walker B Mut.]
B
ss39: Kd = 1.65 μM
Fork (20nt): Kd = 1.35 μM
ds39: Kd = 1.49 μM
5'-overhang (20nt): Kd = 1.18 μM
3'-overhang (20nt): Kd = 1.92 μM
Polarization change (ΔP, mP)
[Csm2-Psy3] (μM)
C
Shu1-Shu2 Dimer
DNA-binding Mut. Shu Complex
ss39
Fork (20nt)
3'-overhang (20nt)
5'-overhang (20nt)
ds39
Polarization change (ΔP, mP)
[Protein] (μM)
D
WT Shu
Walker B Mut.
DNA-binding Mut.
Csm2-Psy3
Buffer
Fluorescence Enhancement (ΔF TNP-ATP) (arbitrary units x 1000)
Wavelength (nm)
E
Walker A Mut. Shu
Walker B Mut. Shu
kDa
LMW
Walker A Mut. Shu Tet.
Walker B Mut. Shu Tet.
Psy3
Shu2
Csm2
Shu1
OD280 (mAu)
Volume (ml)
F
ss39: 5'-(T)20-(random)19-3'*
Kd
WT Shu = 2.76 ± 0.08 μM
Walker A Mut. = 3.60 ± 0.38 μM
Walker B Mut. = 2.81 ± 0.24 μM
Polarization change (ΔP, mP)
[Protein] (μM)

**Figure S1.**

**DNA- and nucleotide-binding with Shu proteins**. (**A**) DNA-binding analysis of WT vs. Shu mutants to dsDNA substrates by Electrophoretic Mobility Shift Assay (EMSA). Protein-DNA complexes were resolved by non-denaturing PAGE: 5 % native polyacrylamide gels. (**B**) DNA-binding analysis of Csm2-Psy3 dimer to ssDNA and dsDNA substrates with different end types by fluorescence polarization assay (FPA). (**C**) FPA DNA-binding analysis of Shu1-Shu2 dimer and DNA binding-deficient Shu complex mutant to ssDNA and dsDNA substrates with different end types. Number in parentheses of substrate names stands for the number of nucleotides in its single-stranded part(s) from the ssDNA/dsDNA junction. (**D**) Fluorescence spectra of TNP-ATP (5 μM) with the Shu proteins, Walker B and DNA binding-deficient mutants (4 μM each). (**E**) Gel filtration and SDS-PAGE of the purified Walker A and Walker B Shu mutants. (**F**) FPA DNA-binding analysis of WT Shu complex and Walker motif mutants with ss39 ssDNA substrate. Dissociation constants ($K_d$) in (**B, F**) were determined by non-linear curve fitting to a one-site binding model from triplicate experiments (details in **Materials and Methods**).

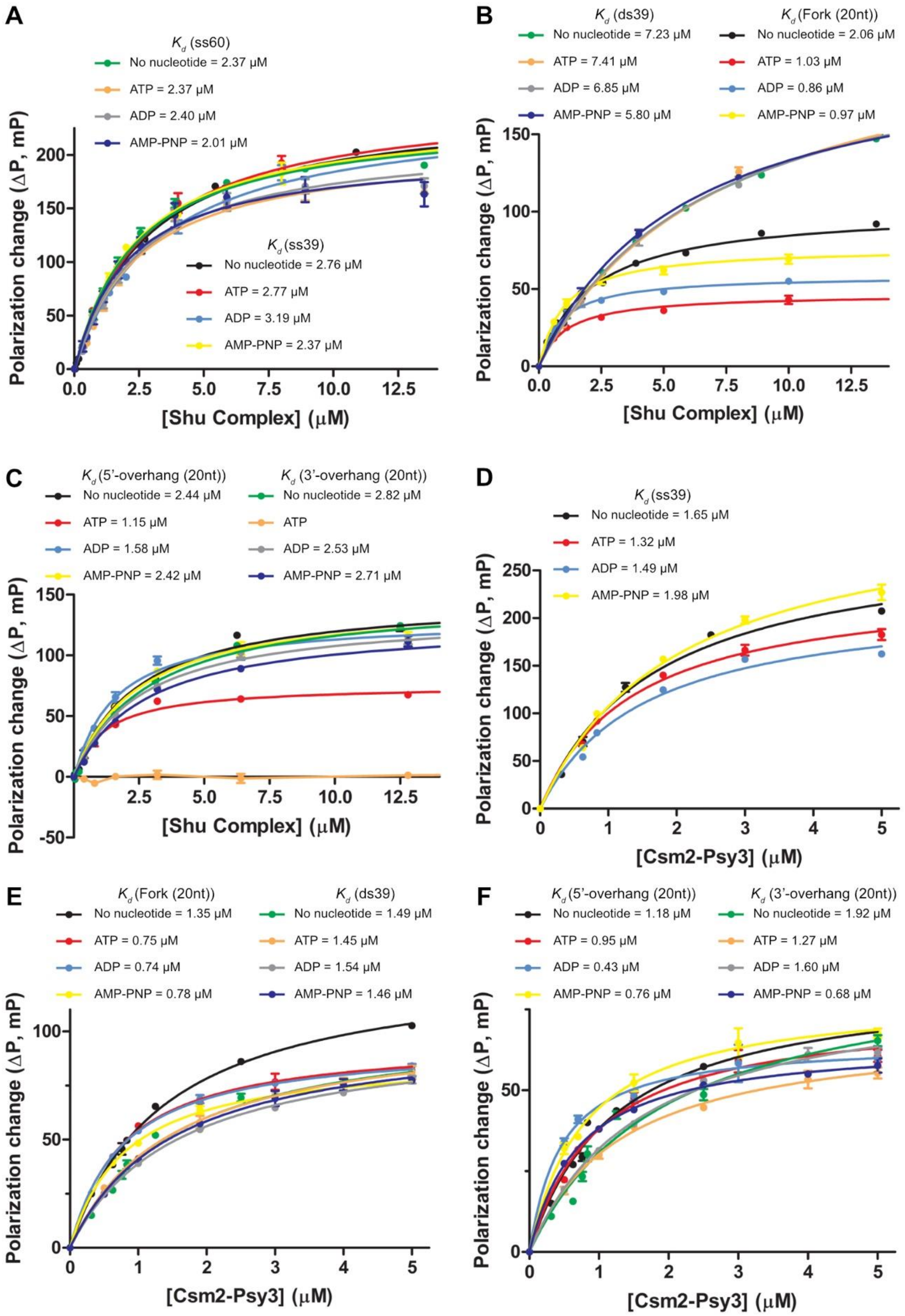
A
$K_d$ (ss60)
No nucleotide = 2.37 μM
ATP = 2.37 μM
ADP = 2.40 μM
AMP-PNP = 2.01 μM
$K_d$ (ss39)
No nucleotide = 2.76 μM
ATP = 2.77 μM
ADP = 3.19 μM
AMP-PNP = 2.37 μM
Polarization change (ΔP, mP)
[Shu Complex] (μM)
B
$K_d$ (ds39)
No nucleotide = 7.23 μM
ATP = 7.41 μM
ADP = 6.85 μM
AMP-PNP = 5.80 μM
$K_d$ (Fork (20nt))
No nucleotide = 2.06 μM
ATP = 1.03 μM
ADP = 0.86 μM
AMP-PNP = 0.97 μM
Polarization change (ΔP, mP)
[Shu Complex] (μM)
C
$K_d$ (5'-overhang (20nt))
No nucleotide = 2.44 μM
ATP = 1.15 μM
ADP = 1.58 μM
AMP-PNP = 2.42 μM
$K_d$ (3'-overhang (20nt))
No nucleotide = 2.82 μM
ATP
ADP = 2.53 μM
AMP-PNP = 2.71 μM
Polarization change (ΔP, mP)
[Shu Complex] (μM)
D
$K_d$ (ss39)
No nucleotide = 1.65 μM
ATP = 1.32 μM
ADP = 1.49 μM
AMP-PNP = 1.98 μM
Polarization change (ΔP, mP)
[Csm2-Psy3] (μM)
E
$K_d$ (Fork (20nt))
No nucleotide = 1.35 μM
ATP = 0.75 μM
ADP = 0.74 μM
AMP-PNP = 0.78 μM
$K_d$ (ds39)
No nucleotide = 1.49 μM
ATP = 1.45 μM
ADP = 1.54 μM
AMP-PNP = 1.46 μM
Polarization change (ΔP, mP)
[Csm2-Psy3] (μM)
F
$K_d$ (5'-overhang (20nt))
No nucleotide = 1.18 μM
ATP = 0.95 μM
ADP = 0.43 μM
AMP-PNP = 0.76 μM
$K_d$ (3'-overhang (20nt))
No nucleotide = 1.92 μM
ATP = 1.27 μM
ADP = 1.60 μM
AMP-PNP = 0.68 μM
Polarization change (ΔP, mP)
[Csm2-Psy3] (μM)

**Figure S2**

**DNA-binding of Shu and Csm2-Psy3 dimer in the presence of different nucleotides.** (**A, B, C**) DNA-binding analysis of Shu complex to (**A**) ssDNA and (**B, C**) dsDNA substrates with different end types, in the presence of different adenine nucleotides by fluorescence polarization assay (FPA). (**D, E, F**) FPA DNA-binding analysis of Csm2-Psy3 dimer to (**D**) ssDNA and (**E, F**) dsDNA substrates with different end types, in the presence of different adenine nucleotides. (A, B, C, D, E, F) Number in parentheses of substrate names stands for the number of nucleotides in its single-stranded part(s) from the ssDNA/dsDNA junction. Increasing concentrations of protein were added to 50 nM 6-FAM fluorescein-labeled DNA substrates. Dissociation constants ($K_d$) were determined by non-linear curve fitting to a one-site binding model from triplicate experiments (details in **Material and Methods**; see **Table S3 & S4** for Shu complex and Csm2-Psy3 data, respectively).

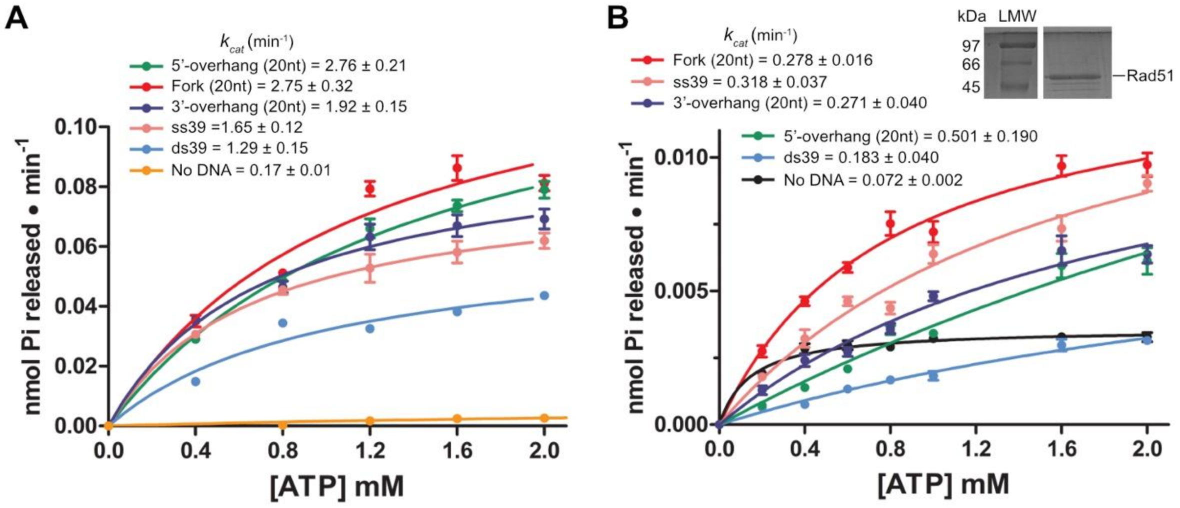


**Figure S3.**

**ATPase activity of Shu complex and Rad51 with different DNA types.** (**A, B**) ATPase activity (90-min reactions) of (**A**) Shu and (**B**) yeast Rad51 (1 μM each) in the presence of DNA (ss39 ssDNA, ds39 blunt-ended dsDNA or dsDNA of the indicated end types: 1 μM each), with increasing ATP concentrations. Data were fitted using the Michaelis–Menten model to determine kinetic parameters with the associated standard error of mean from triplicate experiments (details found in **Table 2** & **Table S5**). Number in parentheses of substrate names stands for the number of nucleotides in its single-stranded part(s) from the ssDNA/dsDNA junction.
